# A pooled genetic approach provides multimodal access to uncharacterized retinal ganglion cell subtypes

**DOI:** 10.64898/2026.08.07.743592

**Authors:** Nina Luong, Olivia Hougham, Joseph Leffler, Benjamin Sivyer, Kevin M Wright

## Abstract

Retinal ganglion cells (RGCs) are the sole output neurons of the retina, responsible for transmitting visual information to the brain. Recent transcriptomic studies have revealed extensive molecular diversity among RGCs that parallels their known morphological and functional heterogeneity. Although large-scale efforts have unified the transcriptomic, morphological, and physiological features for a limited number of RGC subtypes, the majority of molecularly defined types remain poorly characterized. Developing approaches that can be used to identify RGC subtypes in a reliable and reproducible manner is critical for understanding their roles in visual processing. We used a *Mafb^mCherry-2A-Cre^* mouse line to genetically label four uncharacterized RGC subtypes, along with two well-established α-RGC subtypes, and we systematically characterized their molecular markers, central brain targets, dendritic morphologies, and light response properties. Within the population of previously uncharacterized MAFB^+^ RGCs, we identified two OFF-responsive subtypes which we termed “MAFB-midi-OFF” and “MAFB-asymmetric-OFF”, and two ON-OFF-responsive subtypes termed “MAFB-equal bistratified” and “MAFB-unequal bistratified”. Notably, no single molecular, morphological, or functional characteristic was sufficient to distinguish all MAFB^+^ subtypes. Instead, accurate subtype classification emerged only through the integration of multiple complementary features, highlighting the importance of multimodal approaches for defining RGC subtype identity and resolving neuronal diversity within the retina.

## Introduction

Located in the back of the eye, the retina is a thin piece of neural tissue responsible for transmitting a wide range of visual information including motion, color, and changes in luminance to the brain. This is accomplished through parallel circuits formed by synaptic connections between the five neuronal classes in the retina ^1^. Photoreceptors in the outer nuclear layer convert light into electrical signals that are initially transmitted to bipolar cells in the inner nuclear layer, which in turn relay information to retinal ganglion cells (RGCs) in the ganglion cell layer (GCL). This signal is processed and modulated by horizontal cells at the first synapse between photoreceptors and bipolar cell dendrites in the outer plexiform layer, and again by amacrine cells (ACs) at the second synapse between bipolar cell axons and RGC dendrites in the inner plexiform layer (IPL). RGC axons then transmit specific features of the visual scene through the optic nerve to one of more than 50 retinorecipient regions in the brain ^2–4^. Each of these regions is associated with unique visually-guided behaviors, and the location where an RGC axon terminates can provide insights into whether it is responsible for image-forming or non-image-forming visual behaviors ^5^.

A major question that remains unresolved is what constitutes a RGC “subtype”. Classification studies using morphological characteristics including dendritic stratification within the IPL, dendritic field size, branching complexity, and axonal projection patterns have identified at least 35 distinct RGC morphological types “m-types” ^3,4,6,7^. Recent single-cell RNA sequencing (scRNAseq) studies have identified ∼45 molecularly distinct transcriptomic types “t-types” ^8–10^. Unbiased clustering of RGC functional physiological light responses using two-photon calcium imaging identified at least 32 functional types “f-types” ^11^. While multiple RGC subtypes can have shared features within a single modality, a comprehensive classification of a neuronal subtype can be achieved by integrating multimodal m-, t-, and f-type parameters ^12^. Two independent studies used variations of Patch-seq to perform simultaneous physiological recordings, morphological reconstruction, and transcriptomic analysis to classify hundreds of individual RGCs ^13,14^. However, due to the inherent technical difficulties associated with Patch-seq, RGC subtypes that are less abundant or possess small somata are often under-sampled in these studies. Furthermore, this unbiased approach does not allow for the prospective identification of specific RGC subtypes for targeted recordings or manipulations that would be useful for understanding their role in visual circuit development and function.

Transgenic mouse lines that genetically label specific RGC subtypes are a powerful way of overcoming these limitations. This approach has been used for in-depth characterization of multiple RGC types, including α-RGCs, direction-selective RGCs (DSGCs), W3B-RGCs, J-RGCs, F-RGCs, and intrinsically photosensitive RGCs (ipRGCs) ^15–23^. Many of these genetically labeled RGC types can be further split into individual subtypes with distinct morphological and physiological features and a molecular profile that can be matched 1:1 with identified scRNAseq t-types ^8–10,24–26^. Together, these previous studies account for roughly 40% of known RGC t-types, leaving 60% without genetic access. We therefore screened for mouse lines that could potentially be used to genetically target multiple uncharacterized RGC t-types. We identified the transcription factor *Mafb* as a gene with high expression in four RGC t-types (C23, C30, C34, and C45) and lower expression in two other t-types (C39 and C43) ^10^. C23, C30, C34, and C39 are “novel” uncharacterized RGC t-types, while C43 and C45 correspond to α-ON-Sustained and α-OFF-Transient subtypes, respectively. Using a *Mafb^mCherry-2A-Cre^* knock-in mouse line in conjunction with *Cre*-dependent adeno-associated viruses (AAVs), we comprehensively characterized the molecular features, axonal projection patterns, dendritic morphology, and light-response properties of *Mafb*-expressing (MAFB^+^) RGC subtypes. We further integrated these properties to define six putative RGC subtypes: MAFB-asymmetric-OFF-Sustained (C30), MAFB-midi-OFF-Transient (C23/C39), MAFB-equal bistratified-ON-OFF-Transient (C23/C39), MAFB-unequal bistratified-ON-OFF-Sustained (C34), α-OFF-Transient (C45), and α-ON-Sustained (C43).

## Results

### *Mafb* labels an uncharacterized subset of postnatal RGCs

Although several RGC types have been described through morphological and functional approaches, only 18/45 t-types have transgenic mouse lines (*Cre*/*Cre^ER^* or reporter lines) available that provide genetic access for labeling and manipulation studies (**Supplementary Figure 1A**). These transgenic mouse lines rarely label individual RGC subtypes but rather label multiple subtypes. For example, the *Opn4^Cre^*, *Foxp2^Cre^*, and *Kcng4^Cre^* mouse lines label six ipRGCs, four F-RGCs, and four α-RGC subtypes, respectively (**Supplementary Figure 1B**). Using lines that provide genetic access to multiple subtypes simultaneously reduces the number of transgenic lines needed for cell-type classification studies. We mined scRNAseq datasets for candidate genes that were selectively expressed in subsets of uncharacterized RGC t-types, and cross-referenced this list against available transgenic mouse lines that could be used to genetically label RGCs for multimodal analyses (**Figure 1A**) ^8–10^. Across independent RGC atlases at P5 and P56, *Mafb* shows high expression in four t-types: C23, C30, C34, and C45 (α-OFF-Transient) and lower expression in two additional t-types: C39 and C43 (α-ON-Sustained), suggesting that it could serve as a marker for multiple uncharacterized RGC subtypes (**Supplementary Figure 2A, B**) ^10^.

**Figure 1.**
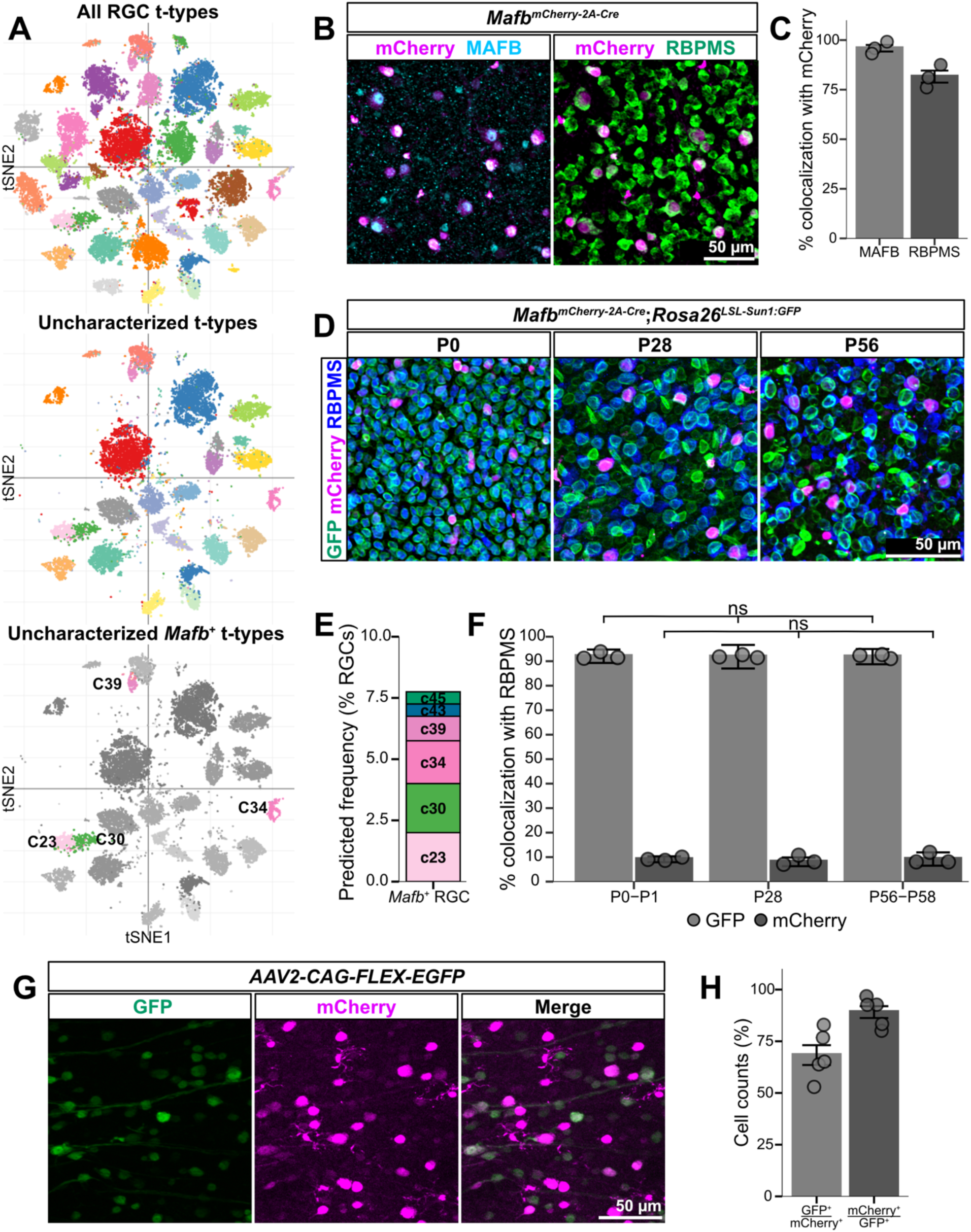
*Mafb^mCherry-2A-Cre^* recapitulates wild type *Mafb* expression and is expressed in retinal ganglion cells. **(A)** Top: tSNE depicting all 45 RGC transcriptomic types (t-types). Middle: tSNE showcasing all RGC t-types that do not have morphology and/or physiology information available. Bottom: tSNE with uncharacterized *Mafb* expressing (*Mafb*^+^) t-types colored. Adapted from Tran et al., 2019. **(B)** *En face* retinas from P21 *Mafb^mCherry-2A-Cre^* mice immunostained with mCherry (magenta), MAFB (cyan) and pan-RGC marker RBPMS (green) to show colocalization. Scale bar = 50 μm. **(C)** Quantification of colocalization with mCherry. Among mCherry^+^ cells, 95.89% ± 1.66% colocalize with MAFB and 81.59% ± 3.04% colocalize with RBPMS. N = 3 animals. **(D)** *Mafb^mCherry-2A-Cre^;Rosa26^LSL-Sun1:GFP^ en face* retinas from postnatal day 0 (P0), P28 and P56 immunostained for GFP (green), mCherry (magenta), and RBPMS (blue). **(E)** Predicted frequency (% RGCs) for *Mafb*^+^ clusters from the Mouse Retina Cell Atlas (MRCA) retina database ^9^. **(F)** Analysis of percent colocalization with RBPMS from P0-P1, P28, and P56-P58 in *Mafb^mCherry-2A-Cre^;Rosa26^LSL-Sun1:GFP^* mice. *Mafb* is expressed in most RGCs throughout development but has stable expression in RGCs postnatally. GFP^+^/RBPMS^+^: P0-1 = 92.02% ± 1.48%; P28 = 91.83% ± 0.54%; P56 = 91.86% ± 0.80%. mCherry^+^/RBPMS^+^: P0-1 = 9.11% ± 0.49%; P28 = 8.07% ± 1.52%; P56 = 9.21% ± 2.25%. N = 3 animals/age. One-way ANOVA with Tukey’s HSD post hoc testing. **(G)** Intravitreal injections (IVT) injections of *AAV2-CAG-FLEX-EGFP* into P28 *Mafb^mCherry-2A-Cre^* mice. Scale bar = 50 μm. **(H)** Percent of cells that are GFP^+^/mCherry^+^ and mCherry^+^/GFP^+^. 68.31% ± 4.79% of mCherry^+^ cells colocalized with GFP, and 89.10% ± 2.87 of GFP^+^ RGCs were mCherry^+^. N = 5 animals. The data are represented as mean ± SEM. Scale bar = 50 μm in **(B)**, **(E)** and **(G)**.

We obtained a *Mafb^mCherry-2A-Cre^* knock-in mouse line that has *mCherry-FLAG-2A-Cre* inserted in-frame downstream of the endogenous *Mafb* locus ^27^. We first validated the *Mafb^mCherry-2A-Cre^* mouse line by immunostaining for endogenous MAFB and mCherry and found near complete colocalization in P21 retinas (**Figure 1B, C**). We observed varied levels of mCherry fluorescence within retinal cells in the *Mafb^mCherry-2A-Cre^* mouse line, which likely reflects the range of endogenous *Mafb* expression (**Supplementary Figure 2B**). We quantified colocalization of mCherry and the pan-RGC marker RBPMS and found that ∼80% of mCherry^+^ cells are RGCs (**Figure 1B, C**), which we refer to as MAFB^+^ RGCs. The remaining ∼20% of the mCherry^+^ population is comprised of ACs and microglia (**Supplementary Figure 2A, C, D**). Based on scRNAseq data, we predicted that MAFB^+^ RGCs should comprise ∼8% of the total RGC population (**Figure 1E**), which matched the observed frequency of mCherry^+^ RGCs from P0 to adulthood (**Figure 1F**).

To examine the dynamics of *Mafb* expression in the developing retina, we bred *Mafb^mCherry-2A-Cre^* mice to the *Rosa26^LSL-Sun1:GFP^* reporter line to genetically label all cells that have ever expressed *Mafb*. Unexpectedly, we found that essentially all RGCs and ACs were GFP^+^ at P0, P28, and P56, whereas only a subset of RGCs, ACs, and microglia were mCherry^+^ (**Figure 1D, F**; **Supplementary Figure 2C, D, E**). These results suggest that *Mafb* is transiently expressed in all RGCs and ACs or their precursors at embryonic ages while expression is maintained in a subset of retinal cells postnatally. The transient widespread nature of *Mafb* expression at embryonic ages precluded combining the *Mafb^mCherry-2A-Cre^* mouse line with a *Cre*-dependent genetic reporter line to selectively label MAFB^+^ RGCs for morphological and functional analyses. To circumvent this limitation, we used intravitreal (IVT) injections of *Cre*-dependent AAV (*AAV2-CAG-FLEX-EGFP*) to selectively label MAFB^+^ RGCs. With this approach, we could control the density of labeled MAFB^+^ RGCs by adjusting the titer of injected AAV. Quantification of *Mafb^mCherry-2A-Cre^* retinas injected with high titer (7×10^11^ vg/mL) *AAV2-CAG-FLEX-EGFP* revealed that 89% of GFP^+^ cells were mCherry^+^, and 68% of mCherry^+^ cells were GFP^+^ (**Figure 1G, H**). These results show that *AAV2-CAG-FLEX-EGFP* can faithfully label MAFB^+^ RGCs with minimal non-specific labeling.

### Combinatorial immunolabeling approach validates six *Mafb*^+^ RGC t-types

To examine MAFB^+^ RGC molecular heterogeneity and to identify secondary markers to distinguish between t-types, we reanalyzed the existing scRNAseq atlas of RGCs in the adult mouse retina (**Supplementary Figure 3A**). We included RGCs from clusters C23, C30, C34, C39, C43 (α-ON-Sustained) and C45 (α-OFF-Transient) which express *Mafb* at varying amounts. It is unclear if there is hidden heterogeneity within populations or if *Mafb*^+^ t-types are closely related to one another. Reclustering of this subset of RGCs confirmed the prediction of six distinct clusters, consistent with the original assignment from the full RGC atlas (**Supplementary Figure 3A**). We next used a one-versus-all approach to define candidate cluster-enriched molecular markers (**Supplementary Figure 3B, C**). Among the top 20 candidate genes, we selected those with antibodies validated for immunohistochemistry. We found that the combination of SPP1, CALB1, and MEIS2 alongside amplification of either endogenous mCherry or viral GFP was sufficient to discriminate four of the six MAFB^+^ RGC t-types (**Figure 2A**). We validated that C43 RGCs (α-ON-Sustained) are SPP1^+^, CALB1^+^, MEIS2^-^whereas C45 RGCs (α-OFF-Transient RGCs) are SPP1^+^, CALB1^-^, MEIS2^+^ (**Figure 2B**, B’ **row 5**, **row 6**). MAFB^+^ RGCs that were SPP1^-^, CALB1^+^, MEIS2^-^ corresponded to C30 (**Figure 2B, B’ row 3**), while MAFB^+^ RGCs that were SPP1^-^, CALB1^-^, MEIS2^+^ (**Figure 2B, B’ row 4**) aligned to C34. We could not identify an immunohistochemical marker that distinguished between C23 and C39, which are both SPP1^-^, CALB1^-^, and MEIS2^-^ (**Figure 2B, B’ row 1**, **row 2**). However, we consistently saw these “GFP^+^ only” somata of different sizes neighboring each other, suggesting the presence of multiple RGC subtype mosaics.

**Figure 2.**
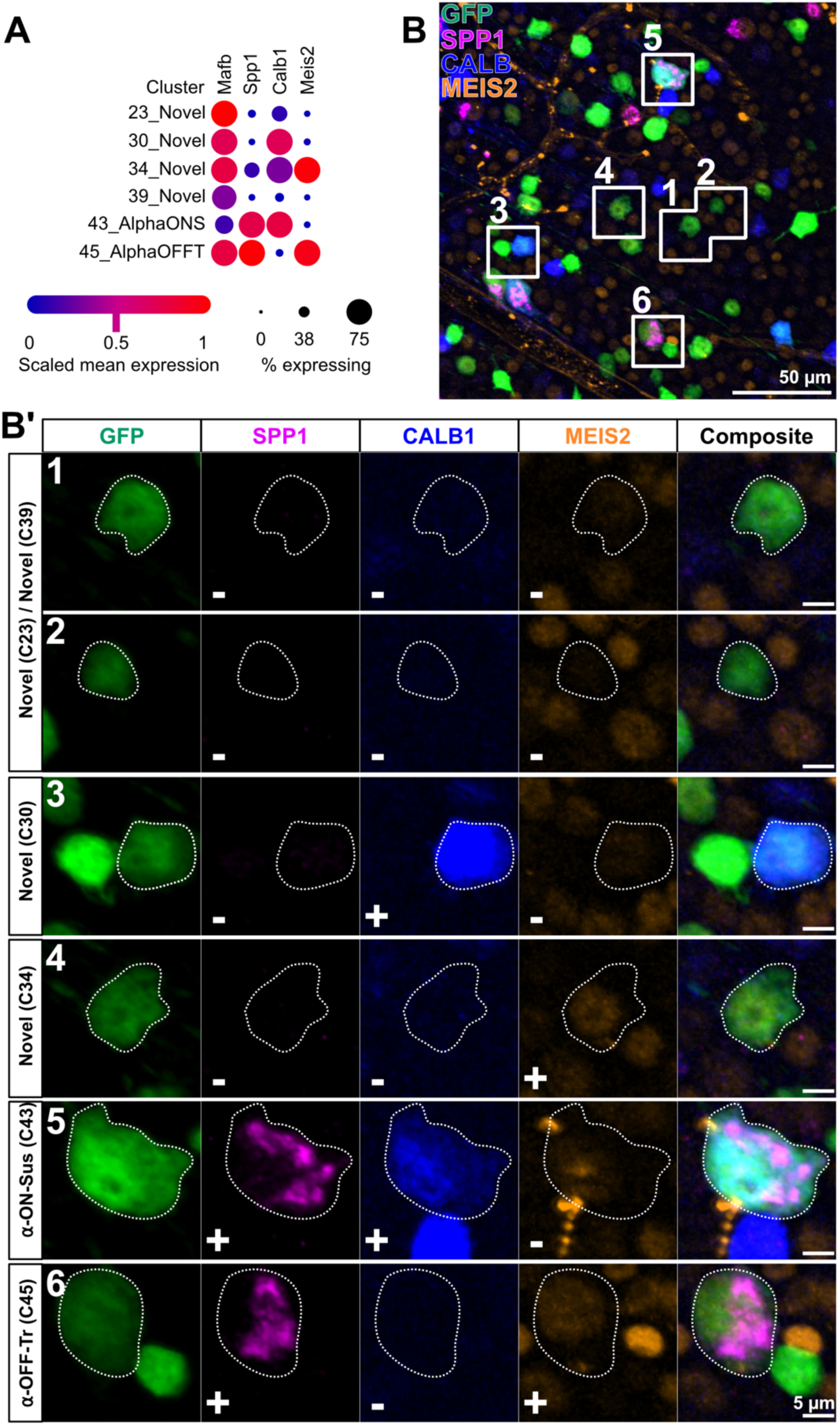
Molecular characterization and validation of *Mafb*^+^ t-types**. (A)** Dot plot showing expression of marker genes across *Mafb*^+^ t-types (Figure adapted from Tran et al., 2019). Marker genes shown include *Spp1*, *Calb1*, and *Meis2* to distinguish α-RGC subtypes from undefined *Mafb*^+^ t-types. **(B)** Immunofluorescence labeling of retinal flatmounts highlighting GFP-labeled MAFB^+^ RGCs and colocalization with molecular markers. Boxed regions **1-6** indicate GFP^+^ cells selected for higher-magnification analysis in panel **B′**. Scale bar = 50 μm. **(B′)** Magnified views of individual GFP^+^ cells corresponding to the numbered regions in panel B. Columns show GFP (green), SPP1 (magenta), CALB1 (blue), MEIS2 (orange), and composite images. Rows correspond to representative cells from clusters C23, C30, C34, C39, α-ON-Sustained (C43), and α-OFF-Transient (C45). Circled regions indicate cell of interest. + / **-** symbol in bottom left region of magnified images indicates if the given marker is present or absent from the cell of interest. Scale bar = 5 μm in composite image, bottom right.

To further validate *Mafb*^+^ t-types, we counterstained with antibodies associated with known subclasses of *Mafb*^+^ and *Mafb*^-^ RGCs: α-RGCs (SMI32/SPP1), F-RGCs (FOXP2), T-RGCs (TBR1), DSGCs (SATB1/SATB2), and T5-RGCs (TUSC5) (**Supplementary Figure 4A**) ^21,28–33^. As anticipated, a subset of MAFB^+^ RGCs (C43, C45) colocalize with the pan-α-RGC markers SMI32 and/or SPP1 (**Supplementary Figure 4B, C, D**). To validate that *Mafb^mCherry-2A-Cre^* labels α-OFF-Transient (C45) and α-ON-Sustained (C43) RGCs, we co-labeled with antibodies for BRN3C and CALB1 respectively and observed a population of α-RGCs triple positive for mCherry, SPP1, and BRN3C (α-OFF-Transient (C45)) and another population triple positive for mCherry, SMI32, and CALB1 (α-ON-Sustained (C43)) (**Supplementary Figure 4C, D**). Based on scRNAseq results, F-mini-OFFs (C4) expresses a negligible amount of *Mafb* (Transcripts per million (TPM) = 0.152) (**Supplementary Figure 2B**) ^10^. We did not detect any colocalization between mCherry and FOXP2, showing that MAFB^+^ RGCs do not overlap with F-RGCs (**Supplementary Figure 4E, J**). We also did not detect any mCherry^+^/TBR1^+^ RGCs, indicating that MAFB^+^ RGCs and T-RGCs are mutually exclusive populations (**Supplementary Figure 4F, J**). SATB1 and SATB2 label a majority of DSGCs, albeit non-exclusively ^29,30^. *Satb1* should be expressed in three of the six *Mafb*^+^ t-types, while *Satb2* is expressed in one (**Supplementary Figure 4A**). In concordance with this, we found that ∼30% of MAFB^+^ RGCs are SATB1^+^, while ∼11% are SATB2^+^ (**Supplementary Figure 4G, H, J**). Four of the *Mafb*^+^ t-types are predicted to express *Tusc5*, and ∼37% of MAFB^+^ RGCs are TUSC5^+^ (**Supplementary Figure 4A, I, J**) ^10^. One RGC subclass lacking an antibody for validation was the W3s, which includes C23 and C30. These two t-types selectively express *Prokr1* and *Postn* respectively, and we confirmed that a subset of MAFB^+^ RGCs are *Prokr1^+^* or *Postn^+^* using RNAscope (**Supplementary Figure 4K, L, M**) ^10^.

Two recent studies examined the spatial distribution of the 45 RGC t-types and show that C23, C30, and C34 have uniform distributions across the retina, whereas C39 exhibits a ventronasal bias and C43/C45 a temporal bias ^25,26^. C23, C30, C34, C39 and C43 have higher nearest-neighbor regularity indices (NNRIs) than matched nulls, indicating spatial regularity ^25,26^. C45 was not analyzed for a NNRI but prior work established that α-OFF-Transient RGCs form an independent mosaic ^21^. Based on our immunohistochemical results and additional information from the spatial transcriptomic mapping studies, the *Mafb^mCherry-2A-Cre^* mouse line labels the six *Mafb*^+^ t-types as predicted from scRNAseq datasets.

### MAFB^+^ RGCs project to image-forming and non-image-forming brain regions

RGCs are specialized to convey information about different aspects of the visual scene to distinct retinorecipient targets in the brain essential for visually evoked behaviors like gaze control (superior colliculus), optokinetic reflex (pretectal nuclei), and circadian rhythms (suprachiasmatic nucleus) ^3–5^. To identify the central targets of MAFB^+^ RGCs we performed either bilateral or unilateral IVT injections of high titer *AAV2-CAG-FLEX-EGFP* in P28-35 *Mafb^mCherry-2A-Cre^* mice and analyzed brains 21 days post-injection. Fluorophore-conjugated cholera toxin B (CTB) was injected either bilateral or unilateral IVT 4 days prior to tissue collection to label all RGC axons and retinorecipient locations (**Figure 3A, B**). We collected serial 100 μm vibratome sections from the entire rostrocaudal axis of the brain and identified retinorecipient regions by the presence of CTB and cross-referenced regions to the Allen Mouse Brain Atlas for assignment ^3,34^. Areas receiving inputs from MAFB^+^ RGCs were identified by colocalization of CTB and GFP.

**Figure 3.**
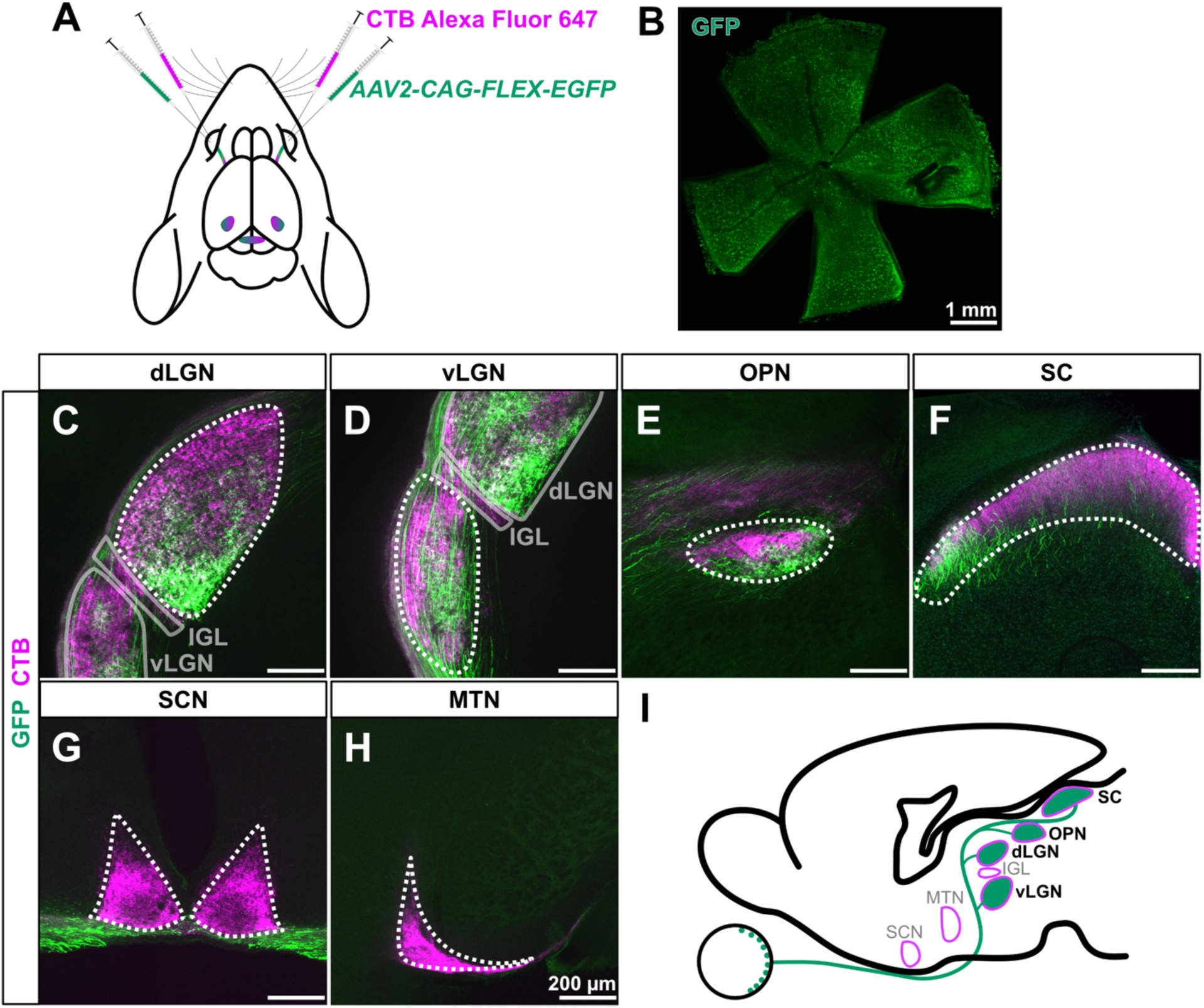
MAFB^+^ RGCs terminate in image-forming and non-image-forming visual areas. **(A)** Schematic of the experimental strategy for labeling and tracing RGC projections. High titer (7×10^11^ vg/mL) *AAV2-CAG-FLEX-EGFP* was injected bilaterally to label MAFB^+^ RGCs and axons with GFP (green). Cholera toxin B (CTB) conjugated to Alexa Fluor 647 (magenta) was used to label all retinal projections (N = 4 animals). **(B)** Example *en face* retina showing widespread GFP-labeled MAFB^+^ RGCs following viral delivery. Scale bar = 1 mm. **(C-H)** Coronal brain sections showing GFP-labeled RGC axons (green) and CTB-labeled retinal projections (magenta). GFP^+^ axons were observed in the **(C)** dLGN, **(D)** vLGN, **(E)** OPN, **(F)** SC. No GFP axons were observed in **(C-D)** IGL **(G)** SCN or **(H)** MTN. Dotted outlines indicate the boundaries of retinorecipient region of interest, whereas semi-transparent solid outlines indicate the boundaries of neighboring retinorecipient regions. Scale bar = 200 μm. **(I)** Schematic of visual targets of MAFB^+^ RGCs. Green denotes observed presence of GFP termination and magenta represents pan-RGC termination.

We observed MAFB^+^ RGC axons terminating in both contralateral and ipsilateral dorsal lateral geniculate nucleus (dLGN), one of the main image-forming retinorecipient locations in mammals that receives innervation from ∼75% of all RGC f-types (**Figure 3C**; **Supplementary Figure 5A**-**A’**, **C**-**C’**) ^35^. The dLGN can be subdivided into shell and core regions, with different f-types preferentially innervating one or the other ^35^. We observed dense innervation of the dLGN core by MAFB^+^ RGCs, and minimal innervation of the shell (**Figure 3C**). The ventral lateral geniculate nucleus (vLGN) was also broadly innervated by contralateral and ipsilateral axons from MAFB^+^ RGCs (**Figure 3D**; **Supplementary Figure 5A**-**A’**, **C**-**C’**). The vLGN supports non-image-forming behaviors like pupillary light reflex, circadian photoentrainment, and light-induced sleep ^36,37^. MAFB^+^ RGCs also sent projections to the contralateral and ipsilateral olivary pretectal nucleus (OPN) core and shell (**Figure 3E**; **Supplementary Figure 5B**-**B’**). The OPN receives input from RGCs encoding luminance information and is the key structure controlling the pupillary light reflex ^38^. The superior colliculus (SC) receives input from 85-90% of all RGCs and is involved in innate responses to visual threats ^5^. We saw broad MAFB^+^ RGC termination throughout all three laminae of the contralateral, but not ipsilateral SC (**Figure 3F**; **Supplementary Figure 5D**-**D’**). We did not observe any MAFB^+^ RGC innervation of the intergeniculate leaflet (IGL), medial terminal nucleus (MTN), or suprachiasmatic nucleus (SCN) (**Figure 3C**, **D**, **G, H**). Overall, the innervation pattern of MAFB^+^ RGCs to dLGN, vLGN, OPN, and SC suggests that they encode both image- and non-image-forming visual information.

### MAFB^+^ RGC are comprised of six m-types

To identify the individual dendritic morphologies of MAFB^+^ RGCs, we performed IVT injections using low titer (1.4×10^10^ vg/mL) *AAV2-CAG-FLEX-EGFP* into *Mafb^mCherry-2A-Cre^* mice and reconstructed and quantified morphometric properties from 103 individual RGCs using the ImageJ/FIJI plug-in Simple Neurite Tracer (**Supplementary Figure 6**) ^39^. We consistently saw several stereotyped RGC m-types that we were able to distinguish from one another using a decision-tree approach based on broad morphometric characteristics including dendritic field size, IPL stratification depth, and symmetry (**Figure 4A**). The α-RGC subtypes (C43 and C45) can be separated from the uncharacterized MAFB^+^ RGC m-types by their large dendritic field (DF) diameter (**Figure 4A, B**; **Figure 5A**). Cells with a DF diameter greater than or equal to 225 μm were classified as α-RGCs and were further subdivided based on IPL stratification depth (α-ON-Sustained (C43): sublamina 5 (S5), α-OFF-Transient (C45): S1) (**Figure 4C**). Cells with smaller DF diameters (less than 225 μm) were then initially separated based on the symmetry of their dendritic arbor. We identified distinct, ventrally oriented MAFB^+^ RGCs with an asymmetric dendritic arbor (**Figure 4B**, **Supplementary Figure 6**). We calculated a symmetry index (SI) and grouped cells with a SI value less than or equal to 0.3 into a single MAFB^+^ m-type which we refer to as “MAFB-asymmetric-OFF” RGCs (**Figure 5B**). RGCs with smaller symmetric dendritic arbors encompass cells with both mono and bistratified dendritic arbors, which we further categorized based on IPL stratification depth (**Figure 4B, C**). Smaller monostratified m-types that stratified in S1 were classified as “MAFB-midi-OFF” (**Figure 4B, C**). Bistratified m-types had a bimodal distribution of dendritic plexus size. One bistratified m-type exhibited dendritic plexi of equal size in S2 and S4, and the other displayed unequal dendritic plexi typically with a larger arbor in S2 and a smaller arbor in S4 (**Figure 4B, C**). To discriminate between the bistratified m-types we calculated the ratio between the dendritic plexi in S4 (DF_1_) and S2 (DF_2_). We next analyzed the bistratifed DF ratio, and if DF_1_ and DF_2_ differed by more than 20%, the cell was categorized as “MAFB-unequal bistratified”. Cells with equivalent dendritic plexi in S4 and S2 were categorized as “equal bistratified”.

**Figure 4.**
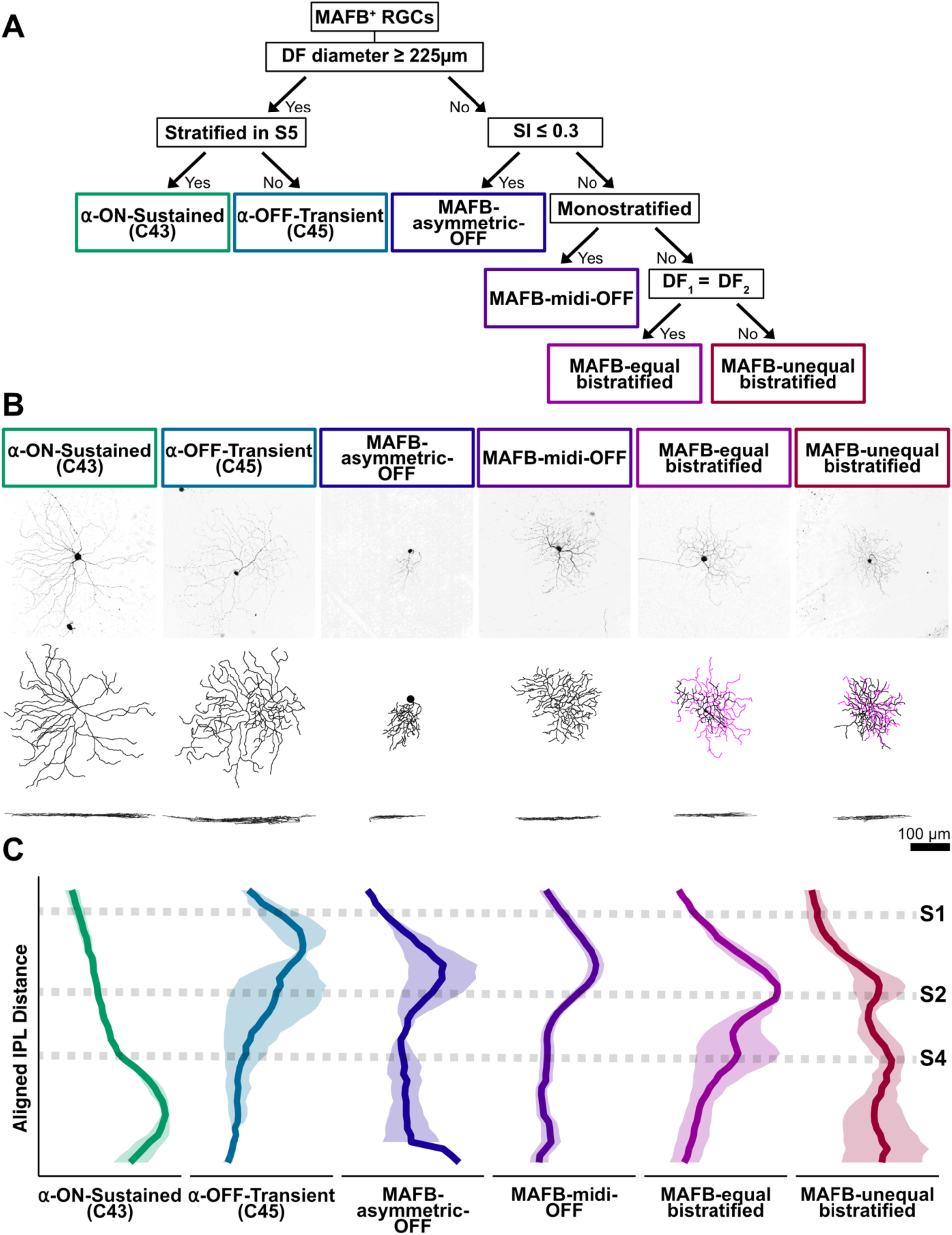
Morphological separation of MAFB^+^ RGCs. **(A)** Decision tree methodology to separate MAFB^+^ RGCs into morphological (m-) types. Cells are first separated based on dendritic field (DF) diameter, followed by stratification in the IPL, symmetry index (SI), dendritic bistratification, and ratio between bistratified dendritic plexi. **(B)** Representative confocal images and dendritic skeletons of MAFB^+^ RGC m-types. Black denotes first dendritic plexus (DF_1_) in S4, and magenta denotes second plexus (DF_2_) in S2 for bistratified cells. Lower image shows z-profile. Scale bar = 100 μm. **(C)** Average IPL stratification of MAFB^+^ RGC. Aligned IPL distance is based on S1, S2, and S4 landmarks using antibodies for tyrosine hydroxylase (TH) and vesicular acetyltransferase (VAChT) respectively. Bold line represents the mean and shaded ribbon indicates SEM. α-ON-Sustained (n = 4 cells), α-OFF-Transient (n = 2 cells), MAFB-asymmetric-OFF (n = 2 cells), MAFB-midi-OFF (n = 19 cells), MAFB-equal bistratified (n = 2 cells), MAFB-unequal bistratified (n = 3 cells), N = 11 animals.

**Figure 5.**
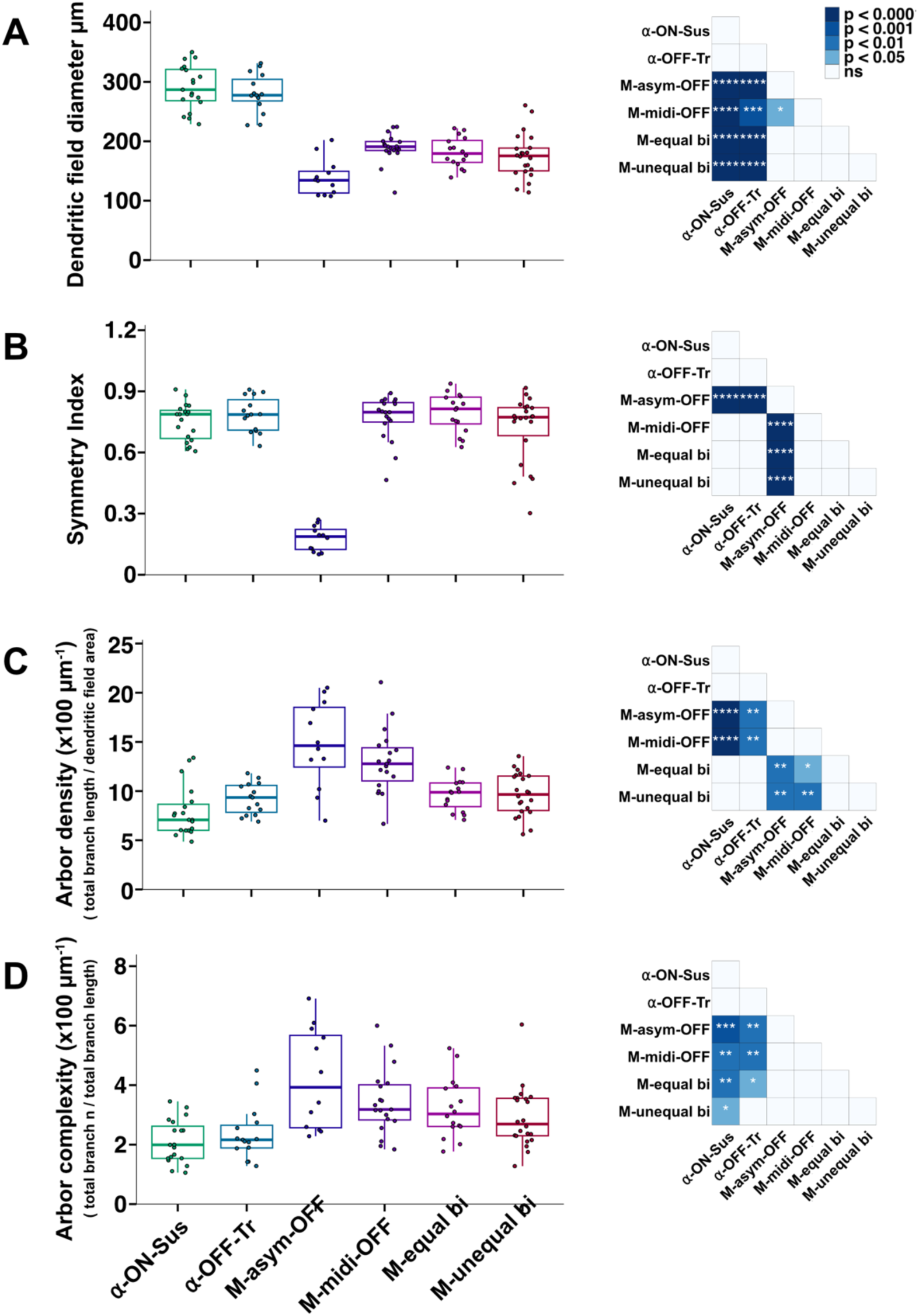
Quantitative comparison of morphological features across MAFB^+^ m-types. **(A-D)** Distribution of **(A)** dendritic field diameter (μm), **(B)** SI, **(C)** arbor density (μm^-1^) ((total branch length / dendritic field area) x 100), and **(D)** arbor complexity (μm^-1^) ((total branch number (n) / total branch length) x 100) across m-types. Each point represents an individual reconstructed cell, with boxplot indicating median and interquartile range. α-ON-Sustained (n = 19 cells), α-OFF-Transient (n = 15 cells), MAFB-asymmetric-OFF (n = 12 cells), MAFB-midi-OFF (n = 19 cells), MAFB-equal bistratified (n = 16 cells), MAFB-unequal bistratified (n = 22 cells); N = 55 animals. A Kruskal-Wallis test, *df* = 5, and post hoc Dunn’s test with Benjamini-Hochberg correction was performed for statistical analysis. Right panel is a pairwise statistical comparison between m-types for each metric displayed as a heatmap. Significance is indicated as: **** = *p* < 0.0001, *** = *p* < 0.001, ** = *p* < 0.01, * = *p* < 0.05, and ns = not significant (*p* ≥ 0.05).

After grouping MAFB^+^ RGC into m-types, we performed statistical analyses of their morphological characteristics (**Figure 5**; **Supplementary Figure 7**, **Table 1**). Across m-types, α-ON-Sustained (C43) and α-OFF-Transient (C45) RGCs exhibited significantly larger DF diameters compared to all other MAFB^+^ m-types (**Figure 5A**). The MAFB-asymmetric-OFF m-type was uniquely characterized by a significantly lower SI relative to other m-types (**Figure 5B**). Significant differences were also observed in arbor density and complexity across m-types. Specifically, the MAFB-asymmetric-OFF and MAFB-midi-OFF m-types displayed higher arbor density than both α-RGC and bistratified m-types and all four uncharacterized MAFB^+^ m-types exhibited greater arbor complexity compared to the two MAFB^+^ α-RGCs (**Figure 5C, D**).

**Table 1.** Summary of morphological characteristics of MAFB^+^ RGCs. Quantified morphological characteristics of MAFB^+^ RGC m-types, including the number of reconstructions (n), furthest z-depth of DF_1_ measured from somal location (μm), furthest z-depth of DF_2_ measured from somal location (μm), soma diameter (μm), DF_1_ diameter (μm), DF_2_ diameter (μm), DF ratio, SI, arbor density (μm^-1^) and arbor complexity (μm^-1^) across reconstructed m-types. Values are displayed as mean (SD).

| m-type | n | z-depth<br>of DF<sub>1</sub><br>($\mu\text{m}$) | z-depth<br>of DF<sub>2</sub><br>($\mu\text{m}$) | Soma<br>diameter<br>($\mu\text{m}$) | DF<sub>1</sub><br>diameter<br>($\mu\text{m}$) | DF<sub>2</sub><br>diameter<br>($\mu\text{m}$) | DF<br>ratio | SI | Density<br>($\mu\text{m}^{-1}$) | Complexity<br>($\mu\text{m}^{-1}$) |
| --- | --- | --- | --- | --- | --- | --- | --- | --- | --- | --- |
| α-ON-Sustained | 19 | 10.8<br>(3.8) | | 15.9<br>(2.4) | 290.4<br>(37) | | | 0.8 (0.1) | 7.9<br>(2.5) | 2.1 (0.7) |
| α-OFF-Transient | 15 | 25.4<br>(9.4) | | 15.4<br>(3.1) | 280.8<br>(32.3) | | | 0.8 (0.1) | 9.2<br>(1.6) | 2.4 (0.9) |
| MAFB-asymmetric-OFF | 12 | 21 (8) |  | 11.7 (2) | 139<br>(30.6) |  |  | 0.2 (0.1) | 14.8<br>(4.4) | 4.2 (1.7) |
| MAFB-midi-OFF | 19 | 20.5<br>(9.1) |  | 12.6<br>(1.7) | 189.3<br>(24.6) |  |  | 0.8 (0.1) | 13<br>(3.2) | 3.4 (1.1) |
| MAFB-equal bistratified | 16 | 13.7<br>(7.9) | 22.5<br>(11.3) | 12.7<br>(1.6) | 181.5<br>(25.3) | 179.9<br>(22.1) | 8.4<br>(4) | 0.8 (0.1) | 9.6<br>(1.6) | 3.2 (1) |
| MAFB-unequal bistratified | 22 | 11.4<br>(4.4) | 22.2<br>(6.2) | 12.5<br>(1.9) | 175.4<br>(37.8) | 190.3<br>(48.2) | 38.9<br>(14.4) | 0.7 (0.2) | 9.6<br>(2.2) | 2.9 (1) |

To determine whether morphometric properties alone could be used to cluster MAFB^+^ RGC m-types, we performed principal component analysis (PCA), k-means clustering, and t-distributed stochastic neighbor embedding (tSNE) using multiple morphometric parameters (**Supplementary Figure 7D**-**G**). Using soma diameter, discrete IPL stratification, bistratifed DF ratio, convex hull, SI, and number of terminal tips to cluster RGC m-types, we identified five distinct clusters (**Supplementary Figure 7D**). Cluster composition showed that the MAFB-midi-OFF and MAFB-asymmetric-OFF m-types each formed distinct clusters, whereas two clusters contained both bistratified m-types comingled, and α-RGC m-types were grouped together within one cluster. Removing the bistratified DF ratio and convex hull, and incorporating DF_1_ and DF_2_ diameter resulted in four clusters, collapsing the two bistratified m-types and the two α-RGC m-types into two clusters (**Supplementary Figure 7E**). This suggests that the selected parameters are insufficient to resolve morphologically similar m-types. Using soma diameter, scaled IPL stratification depth, DF_1_ and DF_2_ diameter, and SI, we identified five distinct clusters (**Supplementary Figure 7F**). These parameters separated the α-RGC m-types into their own clusters, but were still unable to resolve the bistratified m-types from one another. Using discrete IPL stratification, whether a cell was monostratified or bistratified, DF ratio, convex hull, total branch length, and branch length number resulted in six clusters (**Supplementary Figure 7G**). These morphological parameters were sufficient in separating out both the α-RGC m-types and two bistratified m-types. However, upon analysis of each clusters’ composition we noted some intermingling of similar m-types. Overall, morphometric analysis identified six MAFB^+^ RGC m-types using biased criteria, and five to six m-types using unbiased clustering approaches, reflecting the relatively subtle difference between the two bistratified m-types.

### MAFB^+^ RGC m- and t-types show distinct projection patterns

To distinguish which MAFB^+^ RGC subtypes project to the dLGN, OPN, and SC we injected *AAVRetro-CAG-FLEX-EGFP* into each retinorecipient location of *Mafb^mCherry-2A-Cre^* mice to enable retrograde labeling of MAFB^+^ RGCs in the retina (**Figure 6A**). We used a sparse injection strategy to obtain single-cell reconstructions for m-type classification as well as dense labeling combined with the combinatorial antibody method (SPP1, CALB1, MEIS2) for t-type analysis. All MAFB^+^ RGC m-types were represented by retroinjections into the dLGN (**Figure 6A, B**). In contrast, retroinjections into the SC labeled five of the MAFB^+^ RGC m-types except for the MAFB-equal bistratified m-type (**Figure 6A, C**). Retroinjections targeting the OPN labeled the MAFB-asymmetric-OFF, MAFB-unequal bistratified, and both α-RGCs (**Figure 6A, D**). At the molecular level, OPN-projecting cells included C23/C39, C30, C34, C43, and C45 (**Figure 6E**). Notably, most RGCs labeled from dense OPN injections were SPP1^-^, CALB1^-^, and MEIS2^-^, suggesting C23 and/or C39 predominantly innervate the OPN (**Figure 6E**). We were unable to selectively target the vLGN for retroinjections due to its anatomical proximity to the dLGN and IGL, which resulted in viral spread across multiple regions. These results demonstrate that multiple MAFB^+^ RGC m-types project to the dLGN, SC, and OPN.

**Figure 6.**
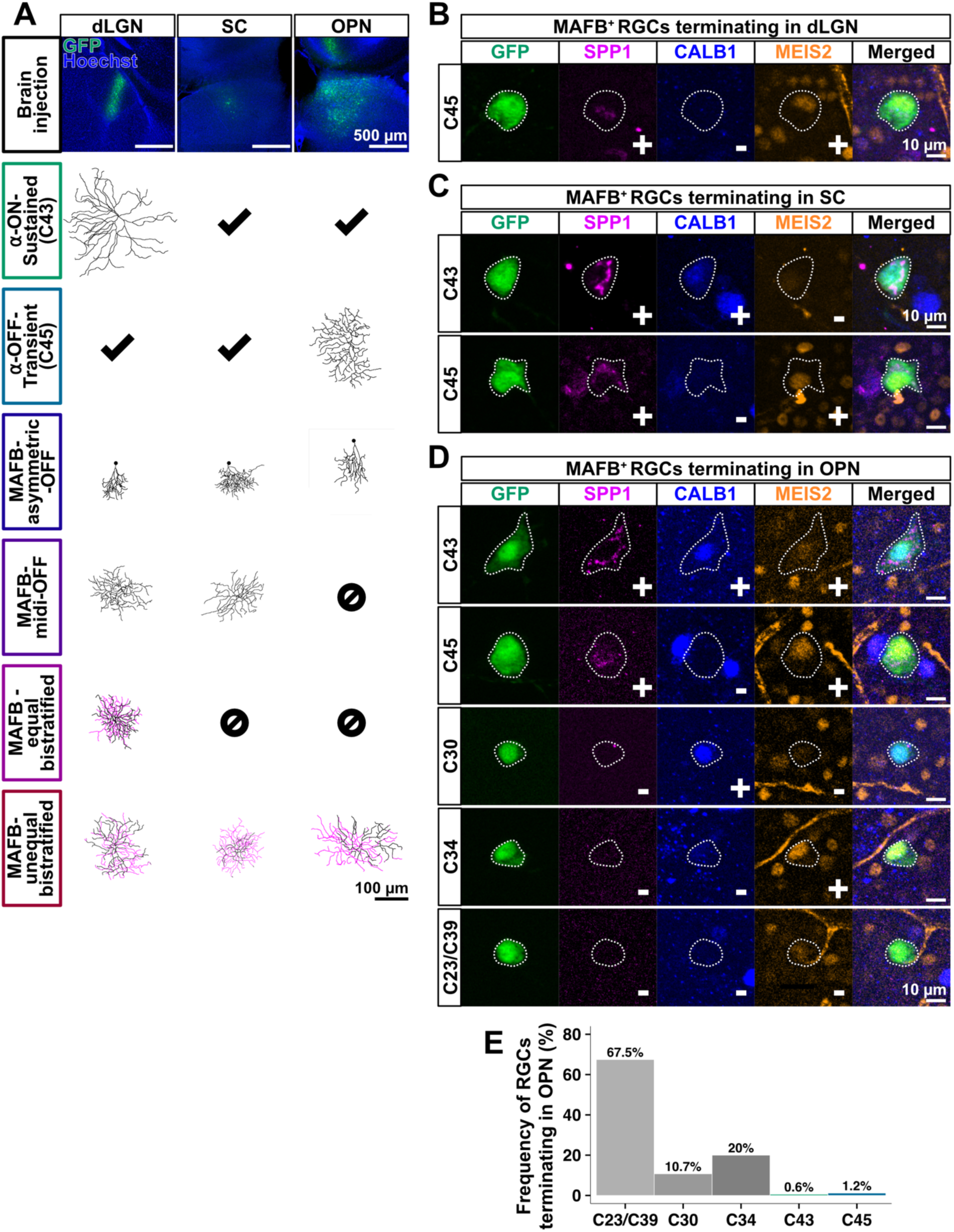
Pairing m-type and t-type to retinorecipient location. **(A)** Example injections into dLGN, SC and OPN using *AAVRetro-CAG-FLEX-EGFP* into P56-65 *Mafb^mCherry-2A-Cre^* mice. Brain slices were immunostained for GFP (green) and Hoechst (blue) to validate injection target. Scale bar = 500 μm. Reconstructed RGCs and their respective m-types are denoted. Check marks indicate RGCs identified through immunolabeling in panels **B-D**. Circle-backslash symbol indicate cells that were not identified from retroinjections. Scale bar = 100 μm. **(B-D)** Retinal flatmounts immunostained for GFP (green), SPP1 (magenta), CALB1 (blue) and MEIS2 (orange) to identify molecular profile of MAFB^+^ RGCs projecting to **(B)** dLGN, **(C)** SC, and **(D)** OPN. Circled regions indicate cell of interest. + / **-** symbol in bottom left region of magnified images indicates if the given marker is present or absent from the cell of interest. Scale bar = 10 μm. **(E)** Relative frequency of MAFB^+^ t-types retrolabeled from the OPN (N = 3 animals). dLGN, N = 8 animals, n = 6 cells reconstructed; SC N = 4 animals, n = 3 cells reconstructed; OPN, N = 3 animals, n = 4 cells reconstructed.

### MAFB^+^ RGCs are comprised of five f-types

To classify MAFB^+^ RGCs based on their functional properties, we used IVT or retrograde AAV brain injections and performed loose-cell attached spike recordings of GFP^+^ neurons. MAFB^+^ RGCs were grouped into f-types post hoc. Based on recordings from 127 RGCs, we identified five distinct MAFB^+^ f-types: ON-Sustained, OFF-Transient, OFF-Sustained, ON-OFF-Transient, and ON-OFF-Sustained (**Figure 7A**-**E**). Baseline firing rates varied across f-types, with ON-Sustained and OFF-Sustained RGCs exhibiting higher baseline activity compared to other f-types (**Figure 8F**). We found that most MAFB^+^ RGC f-types responded optimally to spot diameters of approximately 350-500 μm (**Figure 8A**-**E**). To assess direction selectivity, we presented moving bars and calculated a direction selectivity index (DSI). A subset of ON-OFF-Transient RGCs exhibited DSI values greater than or equal to 0.33 (DSI ≥ 0.33, n = 5 cells), indicative of having a preferred direction (Figure 8H), however the majority had a DSI less than 0.33 (DSI < 0.33, n = 28 cells). To evaluate orientation selectivity, we presented static bars at various angles and calculated an orientation selectivity index (OSI). Similar to direction selectivity, while the majority of MAFB^+^ RGCs were not orientation selective, we occasionally recorded from ON-OFF-Transient MAFB^+^ RGCs that had OSI values greater than or equal to 0.33 (OSI ≥ 0.33, n = 4 cells; OSI < 0.33, n = 25 cells) (**Figure 8I**). We tested contrast sensitivity by presenting a spot that changed in contrast, and we did not observe significant differences in contrast responses across MAFB^+^ RGC f-types (**Figure 8J**).

**Figure 7.**
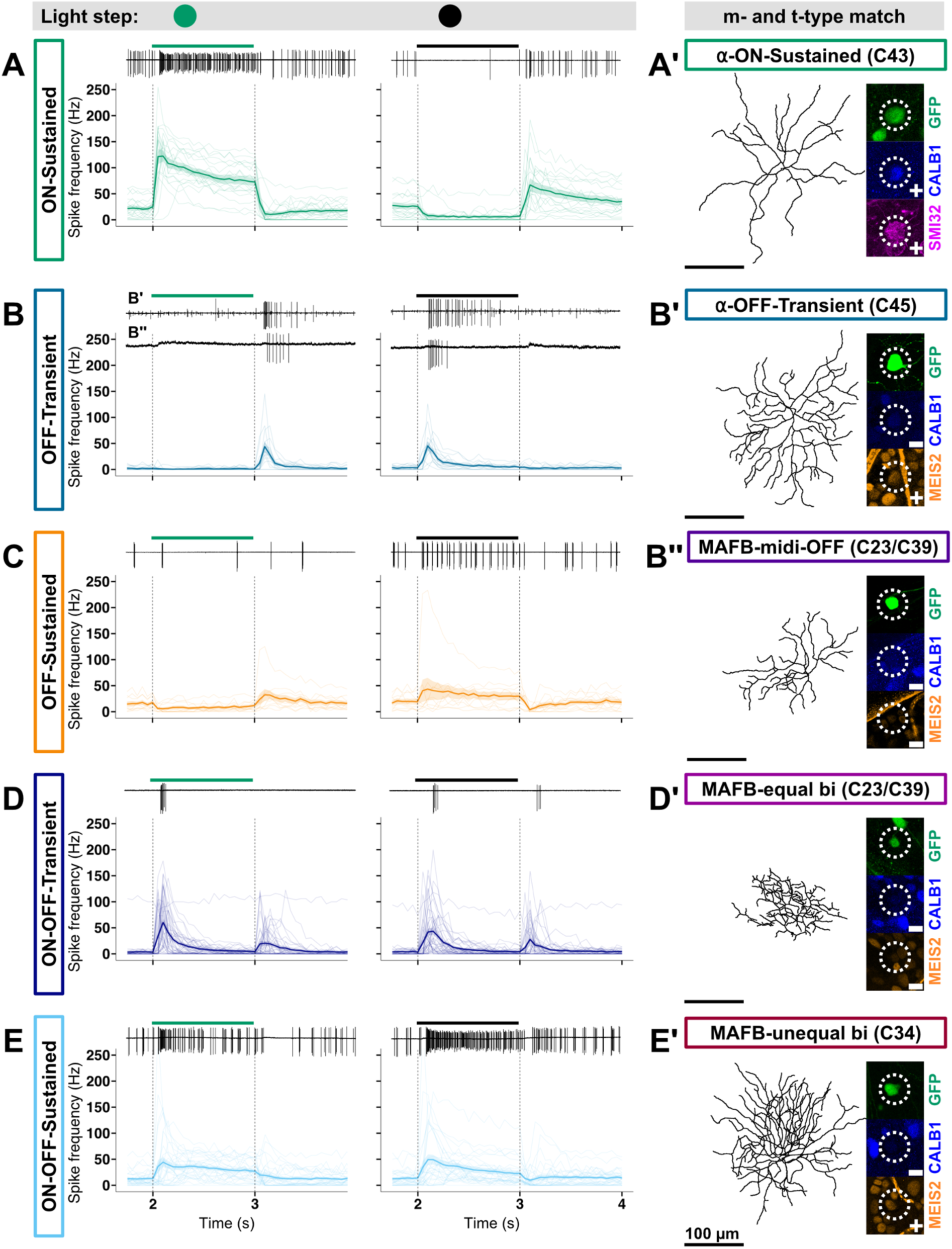
Polarity and kinetics of MAFB^+^ RGCs. **(A-E)** Spike frequency histogram showing responses to an ON light step (green/blue stimuli on grey background; left panel) and OFF light step (black stimuli on grey background; right panel). The mean firing rate (bold line) with individual traces (shaded lines) is plotted for the stimulus duration. The raw traces in the upper left corner of each panel corresponds with the dendritic reconstructions in **A’**, **B’**, **B’’**, **D’**, and **E’**. The bar in each raw trace denotes the presentation of light stimuli. Vertical dashed lines indicate the beginning and end of the stimulus presentation. Cells were classified based on response dynamics into **(A)** ON-Sustained (n = 20 cells), **(B)** OFF-Transient (n = 15 cells) **(C)** OFF-Sustained (n = 11 cells) **(D)** ON-OFF-Transient (n = 46 cells) **(E)** ON-OFF-Sustained (n = 35 cells). N = 63 animals. **(A’-E’)** Molecular and morphological validation of MAFB^+^ RGC t-types. Left panels are representative skeletonized dendritic reconstructions from MAFB^+^ RGC cell fills. Scale bar = 100 μm. Right panels are confocal images showing colocalization of GFP (green), CALB1 (blue), SMI32 (magenta) or MEIS2 (orange) to match m- and t-type. Circled regions indicate cell of interest. + / **-** symbol in bottom left region of magnified images indicates if the given marker is present or absent from the cell of interest.

**Figure 8.**
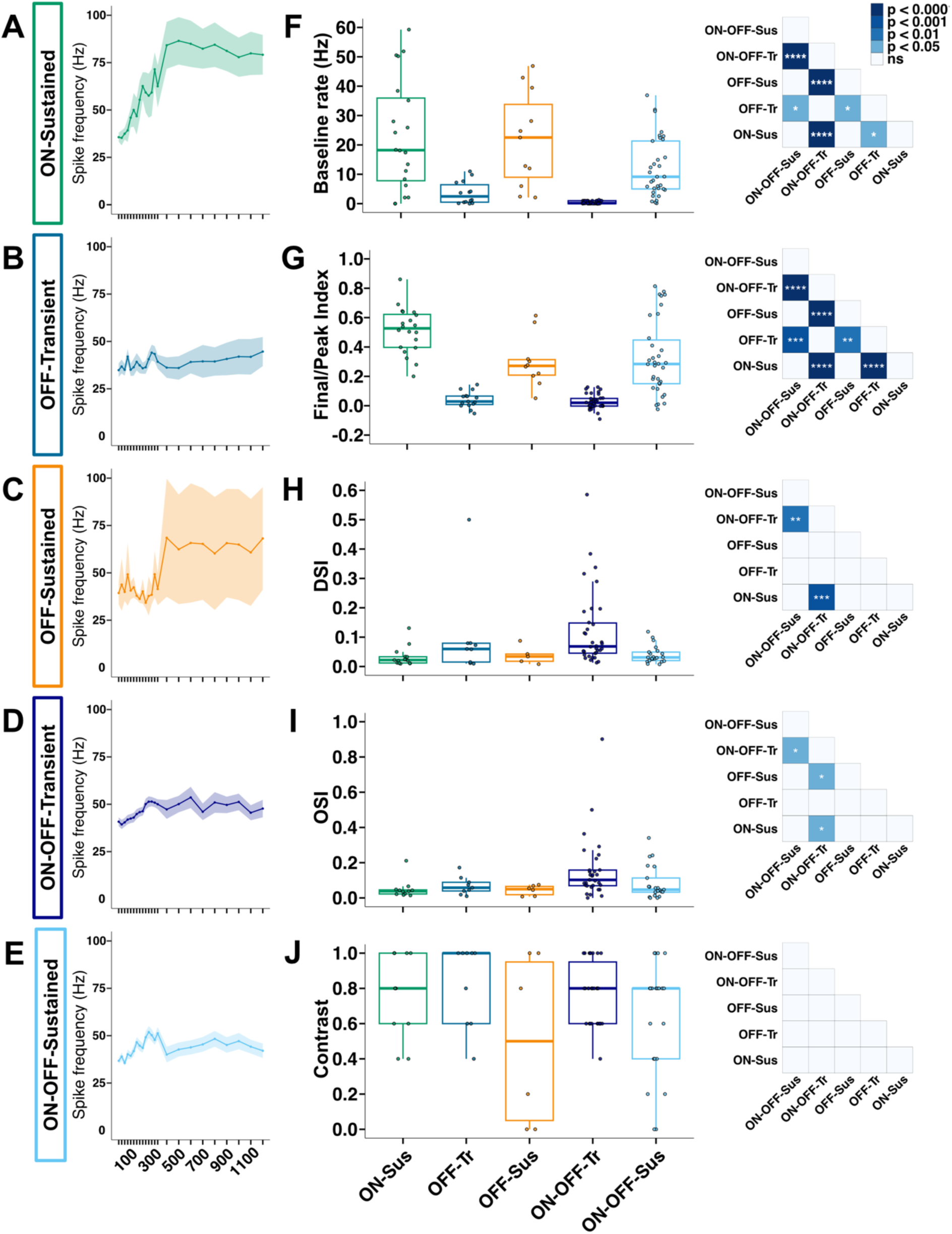
Feature selectivity of MAFB^+^ RGCs. **(A-E)** Average spike frequency (Hz) across spot diameters for MAFB^+^ RGC f-types. Solid lines represent the mean response across cells, with shaded regions indicating SEM. **(F-J)** Quantification of functional properties across f-types including **(F)** Baseline firing rate (Hz), **(G)** Final/Peak (FP) Index, **(H)** Direction Selectivity Index (DSI), **(I)** Orientation Selectivity Index (OSI), and **(J)** Contrast preference. Each plotted point represents an individual cell; boxplots show the median and interquartile range. A Kruskal-Wallis test, *df* = 4, and post hoc pairwise Wilcoxon test with Benjamini-Hochberg correction was performed for statistical analysis. Right panel is a pairwise statistical comparison between f-types for each metric displayed as a heatmap. Significance indicated as: **** = *p* < 0.0001, *** = *p* < 0.001, ** = *p* < 0.01, * = *p* < 0.05, and ns = not significant (*p* ≥ 0.05).

### Unification of MAFB^+^ RGC t-, m-, and f-types

Following the recording session, we performed Neurobiotin fills on a subset of cells to link MAFB^+^ f-types to their corresponding m- and t-types (96 attempted cell fills, 24 cell fills produced complete dendritic labeling for morphological reconstruction). We confirmed that the light response properties of α-ON-Sustained (C43) and α-OFF-Transient (C45) matched previously published two-photon targeted loose-cell attached recordings from genetically identified α-RGCs (**Figure 7A**-**A****’**, **B**-**B’**) ^21^. We recorded from an OFF-Transient RGC that did not match morphologically or molecularly to an α-OFF-Transient RGC (C45); we identified that it was a MAFB-midi-OFF m-type, consistent with C23/C39 (CALB1^-^ and MEIS2^-^) (**Figure 7A**-**A****’**, **B**, **B’’**; **Supplementary Figure 8F**). Bistratified MAFB^+^ RGCs exhibited ON-OFF responses with distinct kinetics. The MAFB-equal bistratified m-type corresponded to the ON-OFF-Transient f-type and was CALB1^-^ and MEIS2^-^, consistent with C23/C39 (**Figure 7D**-**D****’**; **Supplementary Figure 8F**). In contrast, the MAFB-unequal bistratified m-type corresponded to the ON-OFF-Sustained and ON-OFF-Transient f-types and was occasionally CALB1^-^ and MEIS2^+^, consistent with C34 or CALB1^-^ and MEIS2^-^, consistent with C23/C39 (**Figure 7E**-**E****’**; **Supplementary Figure 8F**). Although we consistently recorded from OFF-Sustained RGCs (n = 11 cells), we were unable to recover cell fills to directly confirm their m- and t-type (**Figure 7C**). Based on exclusion, we infer that the OFF-Sustained f-type corresponds to the MAFB-asymmetric-OFF m-type and C30 t-type (CALB1^+^, MEIS2^-^, SPP1^-^) (**Supplementary Figure 8F**).

We next reconstructed dendritic arbors from recorded cells that were successfully filled with Neurobiotin and incorporated these RGCs into our unbiased clustering algorithm to assess if cells grouped by f-type co-clustered based on their m-type (**Supplementary Figure 8A**-**E**). All α-ON-Sustained f-types clustered with α-ON-Sustained m-types and were CALB1^+^ and MEIS2^-^, confirming their t-type identity as C43 (**Supplementary Figure 8A, F**). We obtained two filled α-OFF-Transient f-types, which clustered with α-OFF-Transient m-types. One was CALB1^-^ and MEIS2^+^, consistent with C45, while the other was CALB1^+^ and MEIS2^-^ (C43) (**Supplementary Figure 8B, F**). Similarly, the one filled MAFB-midi-OFF-Transient f-type co-clustered with MAFB-midi-OFF m-types and exhibited a t-type profile aligned with C23/C39 (**Supplementary Figure 8D, F**). Equal bistratified m-types consistently exhibited ON-OFF-Transient f-type responses but showed heterogeneous t-type identities. 1/6 of the MAFB-equal bistratified m-types with ON-OFF-Transient profiles aligned with C34, while 4/6 with C23/C39, and 1/6 could not be assigned to a t-type (**Supplementary Figure 8C, F**). The greatest variability was observed within the MAFB-unequal bistratified m-type, which included both ON-OFF-Transient and ON-OFF-Sustained f-types. Among the MAFB-unequal bistratified cells with ON-OFF-Transient responses, 3/5 aligned with C23/C39, and 2/5 to C34 (**Supplementary Figure 8E, F**). Notably, ON-OFF-Sustained f-types were restricted to the MAFB-unequal bistratified m-type and within this group, 2/5 matched to C23/C39 and 3/5 aligned with C34 (**Supplementary Figure 8E, F**). One MAFB-unequal bistratified cell had an OFF-Sustained response and aligned with C34 (**Supplementary Figure 8E, F**).

### MAFB^+^ RGCs are conserved in non-human primates

While there are 45 RGC t-types found in the mouse retina, primate RGCs have reduced transcriptomic diversity and contain ∼20 t-types ^40^. scRNAseq of foveal and peripheral RGCs in macaque retina found that *Mafb* is expressed in OFF-parasol ganglion cells as well as two foveal RGC clusters (**Supplementary Figure 9A, B**) ^40^. Comparative scRNA-seq analyses reveal that *Mafb* expression is conserved across species; C23 and C30 in the mouse retina map to oRGC6, C34 and α-OFF-Transient RGCs (C45) align with oRGC5, and C39 and two other t-types correspond to oRGC14 ^41^ (**Supplementary Figure 9C**). Consistent with these findings, immunolabeling of macaque retinal tissue with antibodies for SMI32 and MAFB confirmed the presence of MAFB^+^ RGCs (**Supplementary Figure 9D**). While we do not have any information on the morphology or functional properties of primate MAFB^+^ RGCs, their molecular conservation suggests that they have functional importance in visual processing.

## Discussion

One of the major challenges for understanding what neurons do in the visual system is a lack of tools to identify and manipulate specific neuronal subtypes. Prior studies linking m-, f-, and t-types to one another classified hundreds of individual RGCs into unified subtypes, but this approach is labor intensive and low throughput and results in some subtypes being under powered ^13,14^. Genetic tools that allow for the identification of RGC subtypes can facilitate rigorous and reproducible multimodal analysis across labs, yet ∼60% of RGC subtypes lack appropriate mouse lines. In this study, we used the *Mafb^mCherry-2A-Cre^* mouse line to genetically label two well-described RGC subtypes and multiple previously uncharacterized subtypes and unified their molecular (t-type), morphological (m-type), and functional (f-type) properties to comprehensively define each MAFB^+^ RGC subtype (**Figure 9**).

**Figure 9.**
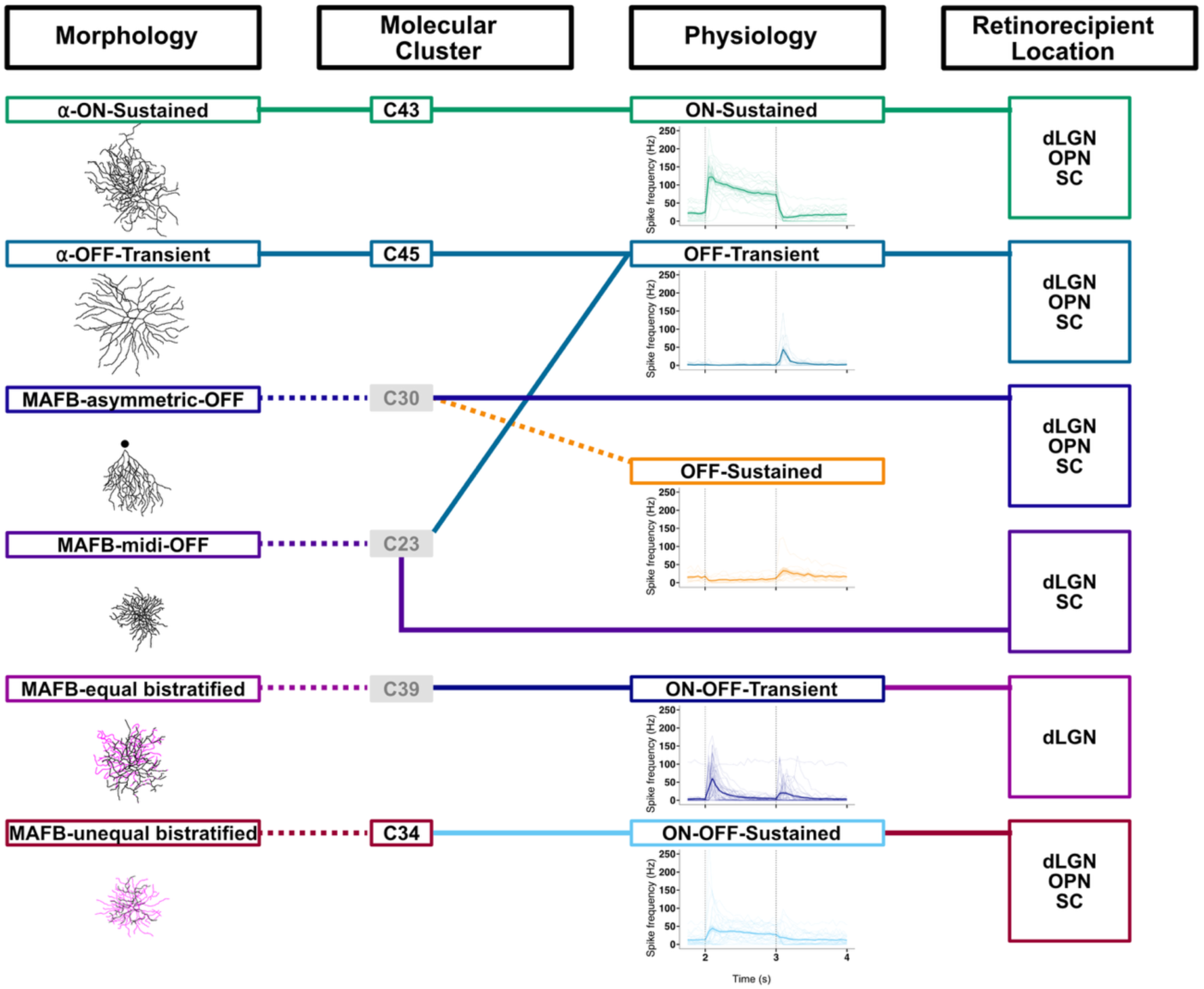
Multimodal classification of MAFB^+^ RGCs. Schematic showing an overview of MAFB^+^ RGC characteristics, and alignment across characteristics. Dotted lines indicate a presumed feature.

The ability to genetically label RGC subtypes has proven instrumental for the classification and in-depth research into the role of α-RGCs, ipRGCs, J-RGCs, W3-RGCs, and DSGCs in vision (**Supplementary Figure 1B**) ^19–21,23,42–53^. The *Mafb^mCherry-2A-Cre^* mouse line adds to these efforts, labeling both known (α-ON-Sustained (C43) and α-OFF-Transient (C45)) and novel RGCs (C23, C30, C34, C39) that are found at relatively low frequencies, with each MAFB^+^ t-type comprising less than 2% for the total RGC population. This relatively low frequency and lack of genetic tools likely explains why these novel subtypes were previously uncharacterized.

Based on combined morphological, molecular, and physiological results, C34 corresponds to the MAFB-unequal bistratified m-type, C30 is most consistent with the MAFB-asymmetric-OFF m-type, and C23/C39 maps to either the MAFB-midi-OFF or the MAFB-equal bistratified m-type. These assignments remain tentative and are limited by the small number of successful cell fills that could be matched to immunolabeling and functional data. Future studies with increased fill success and sample size will be necessary to strengthen these associations. Our findings also highlight that no single modality – morphological, functional, or molecular – is sufficient to unambiguously classify a subtype. Within MAFB^+^ RGCs, we observed substantial overlap in m-type characteristics, f-type properties, as well as variability in t-type identities within the same m-type and f-type groupings, highlighting the need for integrated, multimodal classification of neuronal subtypes.

One of the previously undescribed MAFB^+^ RGCs is a ventrally-oriented asymmetric m-type. This is notable because the mouse retina already contains two genetically-defined, S1-stratfying, ventrally-oriented asymmetric RGCs with similar morphological properties: F-mini-OFFs (C4) and J-RGCs (C5). The DF area of MAFB-asymmetric-OFF RGCs (∼18,000 μm^2^) falls in between that of F-mini-OFFs (∼15,000 μm^2^) and J-RGCs (∼30,000 μm^2^) ^33,49^, and its soma size (∼120 μm^2^) is comparable to F-mini-OFFs (∼100 μm^2^) ^47,54^. Despite their morphometric similarities, these RGC populations have clear molecular distinctions. F-mini-OFFs and J-RGCs belong to the *Foxp2* and *Tbr1* subfamilies, respectively, and do not express *Mafb* (**Supplementary Figure 2B**) ^10^. Conversely none of the predicted MAFB^+^ t-types express *Foxp2* or *Tbr1* by scRNAseq, and we did not detect any colocalization between MAFB^+^ RGCs and FOXP2 or TBR1 supporting the conclusion that the MAFB-asymmetric-OFF RGC is a distinct t-type (**Supplementary Figure 4**) ^10^. Further distinctions between MAFB-asymmetric-OFF, F-mini-OFFs, and J-RGCs emerge when examining axonal projection patterns and functional properties. F-RGCs and J-RGCs preferentially innervate the dLGN shell. In contrast, while MAFB-asymmetric-OFF RGCs project to dLGN based on retrograde AAV labeling, anterograde labeling of all MAFB^+^ RGC axons show minimal innervation of the dLGN shell (**Figure 3C**). F-mini-OFFs project to the OPN while J-RGCs do not, and retroinjections into OPN labeled MAFB-asymmetric-OFF RGCs ^55^. Functionally, both F-mini-OFFs and J-RGCs are OFF-responsive to small spots moving in an upward direction but differ in their kinetics, showing transient versus sustained responses, respectively ^16,33^. In contrast, MAFB-asymmetric-OFF RGCs show OFF-Sustained responses, but do not appear to be direction selective, further distinguishing them from these well-known subtypes (**Figure 7C**). Therefore, although the MAFB-asymmetric-OFF m-type closely resembles F-mini-OFFs and J-RGCs morphologically, they diverge at the level of axonal projection, molecular identity, and light response properties, highlighting the importance of using multiple criteria to define what constitutes a subtype.

Indeed, based on morphometric features alone, it is unlikely that prior studies would have been able to separate MAFB-asymmetric-OFF RGCs from other ventrally-oriented asymmetric types (Eyewire I, Type 2aw) ^7^. Why does the retina contain multiple ventrally oriented asymmetric RGCs? Based on the differences in axonal projection patterns and light response properties, MAFB-asymmetric-OFF RGCs, F-mini-OFFs, and J-RGCs likely function in parallel circuits to encode distinct information. Going forward, large scale connectomic datasets like Eyewire II may provide further insight into this possibility by facilitating the identification of the pre-synaptic inputs onto RGC subtypes ^56^.

MAFB^+^ RGCs include two bistratified subtypes, with relatively similar morphologies but distinct functional properties (transient/sustained firing kinetics) neither of which exhibited strong direction or orientation tuning (**Figure 7D, E**; **Figure 8 H, I**). This is consistent with previous studies identifying bistratified RGCs that are not tuned for motion, direction, or orientation ^57–60^. One example is the RGC labeled by the *Rbp4^Cre^* mouse line (R-cells) that have small somata (soma diameter, 12.9 ± 1.0 μm) and compact dendritic arbors that bistratify in S2 and S4 of the IPL, with one arbor slightly larger and more complex than the other ^61^. R-cells are irradiance-encoding RGCs with sustained firing to light onset with smaller responses to light offset and higher light intensities resulting in reduced spike amplitude. They show weak and inconsistent tuning for direction and orientation. R-cell axons terminate in the dLGN between the external shell and core area, the superficial layers of the SC, and the lateral margin of the vLGN. These morphological and functional characteristics resemble a subset of MAFB-bistratified RGCs, which show a sustained response to both light onset and offset. While the MAFB-bistratified RGCs resemble R-cells morphologically and functionally, it is unclear if they are molecularly the same.

Within the retinofugal pathway there are over 50 target structures that can be grouped into three main circuits: 1) whole-animal physiological states (e.g. circadian rhythm), 2) visually driven reflexive behaviors (e.g. pupillary light reflex, imaging stabilization) or 3) encoding complex visual features. Emerging evidence suggests that RGC f-types responsive to different visual features engage different retinorecipient targets and contribute disproportionately to different behaviors ^5,62^. The functional role of MAFB^+^ RGCs in visual processing remains an outstanding question. Based on their projection patterns to the dLGN, vLGN, OPN, and SC, our data suggest that MAFB^+^ RGCs are likely responsible for controlling visually-driven reflexive behaviors as well encoding complex visual features (**Figure 3C**-**F, I**).

The dLGN is the starting point for higher-order visual representation and perception and can be split into anatomically-distinct shell and core regions that receive eye-specific, retinotopic, and cell-type specific inputs ^35,63–66^. DSGCs preferentially target the shell, while RGC f-types that respond to center-surround stimuli terminate in the dLGN core ^46,47,50,67^. From our anterograde labeling, we see dense innervation of the dLGN core from MAFB^+^ RGCs, in line with prior observations that α-RGCs preferentially target this region (**Figure 3C**) ^68^. In contrast, we observe relatively sparse innervation of the shell, consistent with the lack of direction or orientation tuning in MAFB^+^ RGCs (**Figure 3C**).

The vLGN is responsible for controlling whole-animal physiological states and encoding visual features. The external portion of the vLGN is responsible for supporting non-image-forming visual functions such as eye movement during sleep and features encoding threats and looming dark stimuli ^36,69^. Genetic labeling of ipRGCs and conventional RGCs have shown cell-type specific segregation of the vLGN ^70^. In the rostral portion of the vLGN, projections from all RGC f-types overlap, while the caudal portion of the vLGN is segregated in its innervation by RGC f-types, with ipRGCs terminating only in the core ^71^. DSGC, W3B-, Suppressed-by-Contrast-, and R-RGCs that convey information about direction, motion, contrast, and irradiance innervate the lateral portion of the vLGN ^3,17,19,46,50,57,61,72^. In our study, we observed innervation in the lateral and medial region of the caudal vLGN from MAFB^+^ RGCs (**Figure 3D**). α-ON-Sustained (C43) likely contribute to the innervation seen in the core of the vLGN ^70^. Although we were unable to definitively assign MAFB⁺ RGC subtypes to specific vLGN innervation patterns with retrograde labeling, our data suggest that one or more of the novel MAFB⁺ RGCs contribute to the shell innervation. Given the functional diversity of the vLGN and the limitations in selectively labeling MAFB⁺ projections to this region, the precise role of these cells in vLGN-mediated visual processing remains unclear.

The OPN is a key structure responsible for the pupillary light reflex as it receives input from RGCs encoding luminance information ^73,74^. Retrograde viral labeling and single-cell electrophysiology identified that the OPN receives input from ∼14 RGC f-types including ipRGCs and conventional RGCs, with M1 ipRGCs selectively innervating the shell and M2-M5 ipRGCs terminating in the core region ^22,55,71^. The OPN may also play a role in processing high spatial resolution information based on its connectivity to other visual structures and varied retinal inputs ^13,75,76^. We observed MAFB^+^ RGCs terminating in the core and shell of the OPN (**Figure 3E**). Retrograde labeling revealed that multiple MAFB^+^ RGC t-types including C23/C39, C30, and C34 contribute to this innervation (**Figure 6A**, **D**, **E**). Although we were not able to resolve subtype-specific targeting within the core versus shell region, the broad innervation of the OPN by MAFB^+^ RGCs suggests they may participate in both visually driven reflexive behaviors and encoding complex visual information.

About 90% of all mouse RGCs project to the SC, which integrates head and eye movements and initiates defensive behaviors in response to visual stimuli that represent potential threats like expanding dark stimuli ^3,77,78^. RGC f-types that are sensitive to direction, motion, and orientation tend to innervate the upper layers of the SC, while RGCs sensitive to center-surround stimuli, like α-RGCs, terminate in the lower part of the SC ^43,70,79–81^. Based on the widespread innervation we observed across all layers of the SC, MAFB^+^ RGCs may contain f-types that are selective for motion that our stimulus set could not detect (**Figure 3F**).

scRNAseq datasets provide an ideal starting point for identifying new genetic tools to label and manipulate neuronal subtypes of interest. However, one limitation is that there are few transgenic lines that provide access to just a single RGC subtype. In addition to *Mafb^mCherry-2A-Cre^*, several other transgenic lines mark multiple subtypes that are related to each other for example, ipRGCs in the *OPN4^Cre^* mouse and α-RGCs in the *Kcng4^Cre^* mouse ^20,21^. While lines that label multiple putative subtypes are ideal for characterizing and unifying m-, t-, and f-properties, they are less ideal for understanding what specific RGC subtypes do with respect to visual behaviors. For these studies, more sophisticated intersectional genetic approaches will be needed to target single subtypes. For example, pairing a subclass-specific driver like *Mafb^mCherry-2A-Cre^* with a *Flp* mouse could be used to gain single subtype specificity. When cross referencing RGC scRNAseq atlases, we identified secondary genes enriched in each MAFB^+^ RGC cluster that could be used in this approach: C23 (*Kcnip4*, *Nfic*), C30 (*Prdm16*), C34 (*Tshz3*), C39 (*Nfia*), C43 (*Esrrg*, *Irx1*) and C45 (*Kcnip2*) ^8,10^.

Another outstanding question is whether *Mafb* plays a role in the development or function of MAFB^+^ RGCs. The 45 RGC t-types in mouse can be subdivided into eight subclasses based on the selective expression of a specific transcription factor (*Eomes, Tfap2d, Bnc2, Mafb, Foxp2, Irx3, Tbr1, Neurod2*) ^82^. *Eomes* is selectively expressed in ipRGCs, where it is essential for their development and function ^83–86^. TBR1^+^ RGC subtypes are all OFF-stratifying, and genetic deletion of *Tbr1* results in the abnormal dendrite development and mis-stratification of these RGC subtypes ^87,88^. FOXP2^+^ RGCs include four closely related subtypes with shared morphological features (F-mini-ONs, F-midi-ONs, F-mini-OFFs, F-midi-OFFs), although the function of *Foxp2* in these cells remains unknown. Based on our results, there are no obvious morphological or functional properties shared by all MAFB^+^ RGC subtypes.

Transcriptomic analysis shows that among the MAFB^+^ RGC subtypes, α-ON-Sustained (C43) shows the strongest enrichment for ON-associated gene expression. OFF-associated genes are most enriched in C34 and α-OFF-Transient (C45), with lower expression in C23 and C30. In contrast, C39 exhibits gene expression patterns more consistent with ON-OFF RGCs ^14^. Testing whether *Mafb* plays a role in the development or function of MAFB^+^ RGCs will require different genetic approaches, as *Mafb^-/-^* mice die at birth due to respiratory dysfunction and the *Mafb^mCherry-2A-Cre^* line retains a functional copy of *Mafb* ^89^.

By integrating morphology, physiology, and molecular identity, this work establishes a unified framework for defining MAFB⁺ RGC subtypes. More broadly, our findings highlight the importance of multimodal approaches for resolving neuronal diversity and provide a foundation for understanding how these conserved RGCs contribute to visual processing.

## Methods

### Key resources table

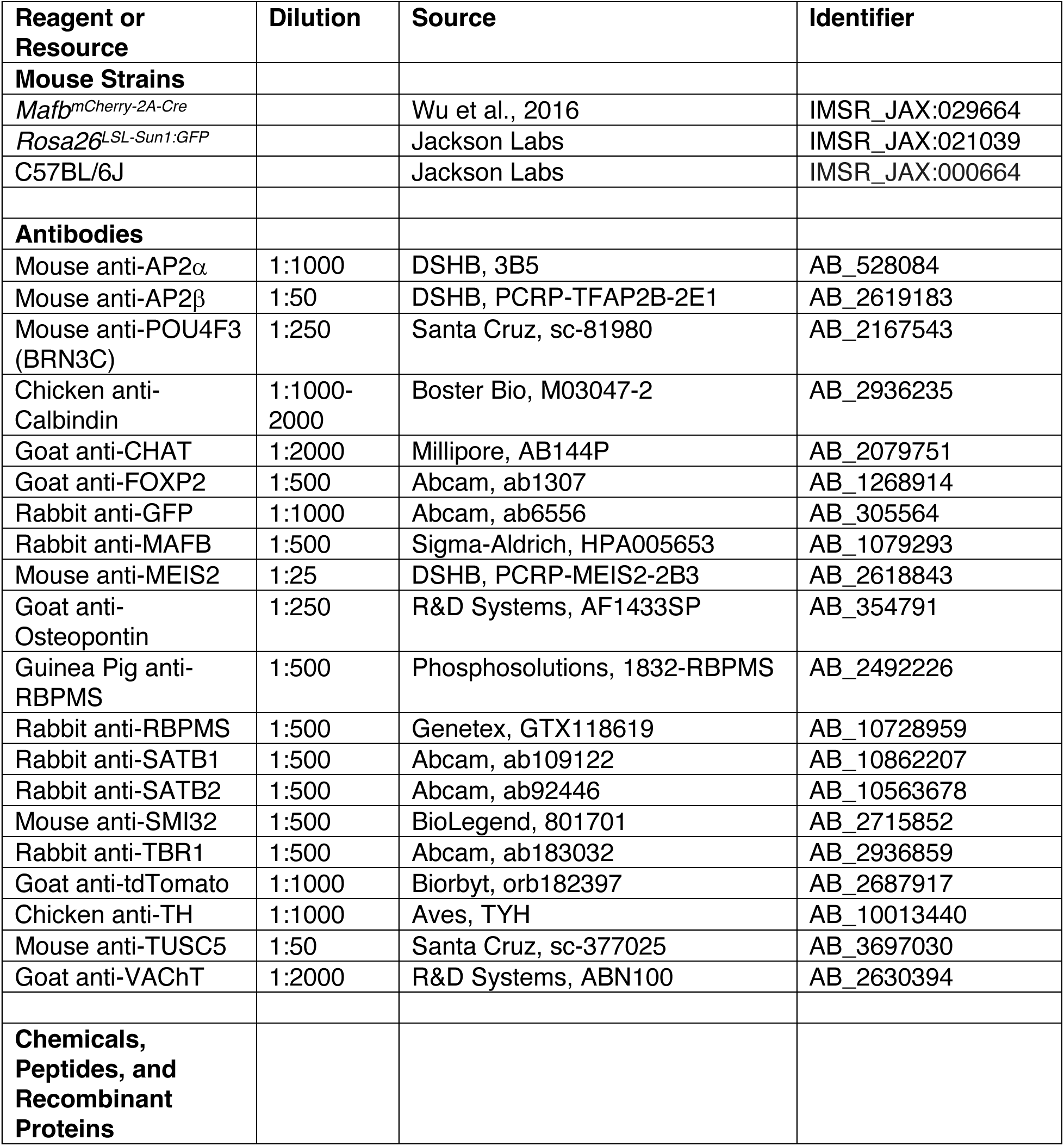
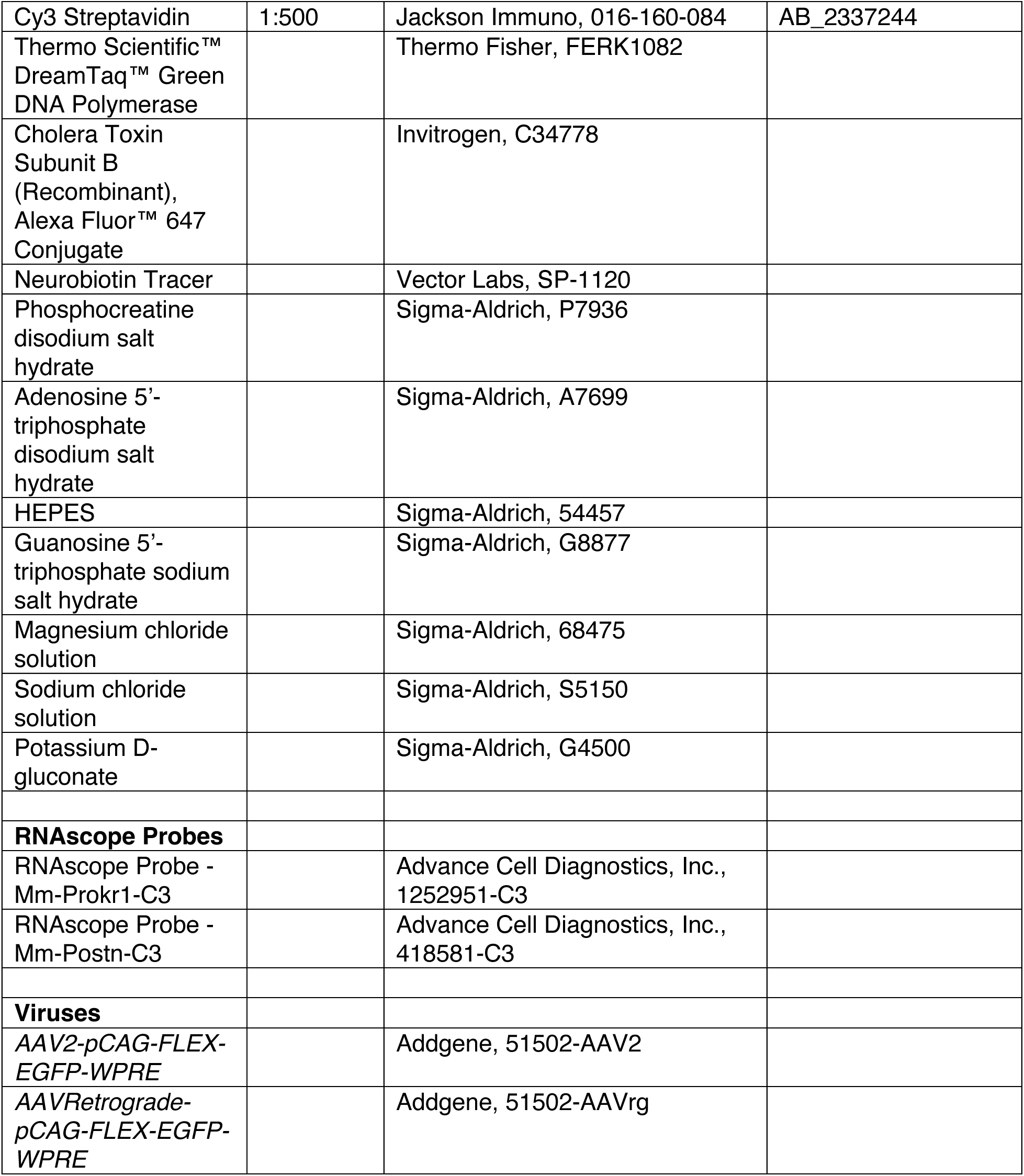

### Animal husbandry

Animals were housed and cared for by the Department of Comparative Medicine (DCM) at Oregon Health & Science University (OHSU), an AAALAC-accredited institution. Animal care, usage, and experimental procedures were approved by OHSU’s Institutional Animal Care and Use Committee (Protocols: IP00000096 and IP00000539), in compliance with the NIH Guide for the Care and Use of Laboratory Animals, and were provided with 24-hour veterinary care. Animal facilities are temperature and humidity regulated and maintained on a 12-hour light-dark cycle. Animals were group-housed. Food and water were provided *ad libitum*. Mice were used between the ages of P0 and P120 (indicated in the text or figure legend). All animals were euthanized by administration of CO_2_ or isoflurane followed by decapitation, cervical dislocation, or exsanguination.

### Mouse strain and genotyping

The day of birth was designated postnatal day 0 (P0). The ages of mice used for specific experiments are indicated in the figures and legends. *Mafb^mCherry-2A-Cre^* mice were obtained from Jackson Laboratory. Male homozygous *Mafb^mCherry-2A-Cre^* mice were bred to C57BL/6J females to generate heterozygous *Mafb^mCherry-2A-Cre^* mice used for all experiments. Genomic DNA extracted from toe or tail samples using the HotSHOT method was used to genotype animals ^90^. The following primers were used to genotype *Mafb^mCherry-2A-Cre^* mice: AGAACGAGAAGACGCAGCTC (common forward), GGCGCAGAATAGGGAGTCT (WT reverse), and GCGCATGAACTCCTTGATGA (mutant reverse). *Rosa26^LSL-Sun1:GFP^* mice were genotyped using a separate reaction and the following primers: AAGGGAGCTGCAGTGGAGTA (WT forward), CAGGACAACGCCCACACA (WT reverse), ACACTTGCCTCTACCGGTTC (mutant forward) and CTGAACTTGTGGCCGTTTAC (mutant reverse). Animals of both sexes were used in this study and data from both sexes were combined for analysis.

### Intravitreal injections for labeling MAFB^+^ RGCs

Adult animals (P28-P35) were anesthetized using isoflurane at an initial flow rate of 3%, and maintained at a flow rate of 1.5%. After being deeply anesthetized, animals were placed on a Kopf stereotaxic injection rig (Model 1900 Stereotaxic Alignment system). 0.5% proparacaine hydrochloride ophthalmic solution (Sandoz, NDC no. 61314-016-01) was applied as a topical anesthetic to the eye, followed by 1% tropicamide ophthalmic solution (Somerset Pharma, NDC no. 70069-121-01) for dilation. Systane Lubricant Eye Gel (Alcon, NDC no. 0065-0474-01) was added to both eyes, and a micro vessel clamp was used to push the globe outwards. For sparse labeling of RGCs, a beveled glass pipette was loaded with *AAV2-CAG-FLEX-EGFP* (Addgene, Plasmid No. 51502, Cat. No. 51502-AAV2) at a titer of 1.4×10^10^ vg/mL. 1μL was injected into each eye using a Nanoinject III (Drummond Scientific Company, Cat. No. 3-000-207). For dense labeling of RGCs and mapping retinorecipient locations, 1-2 μL of *AAV2-CAG-FLEX-EGFP* at a titer of 7×10^11^ vg/mL was injected into each eye. To label all retinorecipient locations, 1-2 μL of 647-conjugated CTB was IVT injected 4-days prior to euthanizing the animal. For all IVT injections, the glass pipette was left in place for 1 minute following all injections. Animals were euthanized 7-21 days post AAV-injection.

### Retrograde labeling of MAFB^+^ RGC axons

Adult animals (P56-P65) were anesthetized using isoflurane at an initial flow rate of 3%, and maintained at a flow rate of 1.5%. After d anesthetization, animals were placed on a Kopf stereotaxic injection rig. Ear bars were adjusted to hold skull in place. Puralube ophthalmic ointment (Chewy, Cat. No. 83592) was added to both eyes. The skull was leveled, and the following coordinates were scaled to a bregma to lambda distance of 3.8 mm: dLGN (ML: ± 2.25, AP: −2.2, Z: −3.25, −3.05, −2.85, −2.65, −2.45, −2.25, −2.05), vLGN (ML: ± 2.55, AP: −2.3, Z: −3.0), SC (ML: ± 0.6, AP: −3.53, Z: −1.07), and OPN (ML: ± 0.6, AP: −2.5, Z: −2.37). A small hole in the skull was drilled above the injection region. The dura was removed/punctured using a 32G needle. Retinorecipient brain regions were injected with undiluted *AAVRetro-CAG-FLEX-EGFP (*Addgene, Plasmid Cat. No. 51502, Cat. No. 51502-AAVrg) at a titer of 7×10^12^ vg/mL through a beveled glass pipette using a Drummond Nanoinject III. The needle was left in the injection target region for 5 minutes before injecting 100-250 nL of virus at a speed of 2 nL/s. The needle was allowed to remain in place for 5 minutes after injection to prevent leakage. Animals were euthanized 21 days post brain injection.

### *Ex vivo* retina preparation

Adult animals IVT injected with high titer (7×10^11^ vg/mL) *AAV2-CAG-FLEX-EGFP* were used for electrophysiology experiments. Mice of either sex were dark adapted for a minimum of 1 hour and anesthetized with isoflurane followed by cervical dislocation. Dissections were performed under infrared illumination (940 nm) with night-vision goggles and an additional infrared light source. Eyes were enucleated, punctured with a 32G needle to relieve pressure and the cornea, lens, and vitreous were removed before the eye cup was placed in carbogenated (95% O_2_, 5% CO_2_) bicarbonate-buffered AMES solution (USBiological Life Sciences, Cat. No. A1372) at 23-24°C. The left and right eye were kept separate and cardinal directions were defined using scleral landmarks. Retinas were then cut into ventral and dorsal hemispheres. Hemisectioned retinas were mounted on an anodisc membrane (Whatman, Cat. No. WHA68097023) with the ganglion cell layer up and placed in a recording chamber and held down via a slice anchor. The recording dish with the oriented retina was placed on the recording rig, and perfused with warmed carbogenated AMES (34-36°C) at a flow rate of 2 mL/min.

### Electrophysiology recordings

Single cell loose cell attach recordings were performed using borosilicate glass patch electrodes (2-8 MΩ) filled with AMES solution. Recording data was collected using a HEKA EPC-10 USB amplifier and Patchmaster software (2.92). Cells were visualized through a 10x and 60x water-immersion objective and differential interference contrast imaging. Fluorescent RGCs were targeted using a brief 554 nm exposure (< 100ms) and a high-sensitivity camera (Andor technologies Model No. DU-888E-COO-#BV). Visual stimuli were presented using a Texas Instruments DLP4500 LightCrafter projector and custom software (pyStim; https://github.com/SivyerLab/pyStim) at a frame rate of 60 Hz. Visual stimuli were generated on a 912×1140 pixel digital projector using a 4-channel LED array with 405 nm (UV), 470 nm (blue), 530 nm (Green), and 590 nm (Amber) with LED channels focused on the photoreceptor layer. All visual stimuli were presented using a grey background (4.45 × 10^13^ photons/cm^2^·s) and repeated 2-3 times. Depending on recording location and color preference, either green or blue stimuli were used to evoke the strongest response. Photon flux was attenuated to desired levels using neutral density filters (Thorlabs).

The RF center of individual RGCs was determined by identifying the maximal firing rate to 4 Hz horizontal and vertical bars (100 x 500 μm) flashed at 100 μm intervals at 13 locations along each axis. The following stimuli were presented at the RF center. To identify spot size preference, diameters ranging from 50-1200 μm were presented at pseudorandom. The diameter corresponding with the maximal firing rate was used for identifying polarity. Polarity and kinetics were defined using a generic color (green or blue) and black light step with the preferred spot size for 1 second stimulus on, followed by 1 second of stimulus off. Contrast preference was determined by presenting the preferred spot size at six contrast levels (α = 0-1, Δ = 0.2 increments) in a pseudorandom sequence. Orientation selectivity was defined using bars (800 x 50 μm) flashed for 1 second at 12 different angles, spaced 30° apart. Direction preference was determined by moving bar stimuli consisting of a rectangular bar (200 x 800 μm) passing through RF center at a speed of 1000 μm/s across 12 different angles spaced every 30°. For all visual stimuli either green, blue, or black stimuli were used based on which stimulus evoked the largest response.

After recording, RGCs were filled using loose-cell attached electroporation with patch pipettes filled with 2% Neurobiotin tracer (Vector Laboratories, Cat. No. SP-1120) diluted in intracellular solution composed of the following (mM): 135 Potassium Gluconate, 7 NaCl, 10 HEPES, 10 Phosphocreatine, 2 Na2-ATP, 0.3 Na-GTP, 2 MgCl_2_ at ∼280-300 mOsm, pH ∼7.2. Hemisectioned retinas were washed with perfused warm AMES solution for 25 minutes before being fixed with 4% EM-grade paraformaldehyde (PFA) (Electron Microscopy Sciences, Cat. No. 15700) for 25 minutes at room temperature. EM-grade PFA was purchased at 16% stock solution and diluted in 0.1M PBS to obtain a final concentration of 4% PFA. Tissue was washed 3 x 10 minutes with 0.1M PBS. Following fixation and washes, retinas were prepped for immunohistochemistry following the protocol for flatmounts below.

### Tissue preparation and immunohistochemistry

For experiments where both retina and brain tissue were collected, animals were deeply anesthetized using CO_2_ and intracardially perfused with 25 mL of 0.1M PBS followed by 25 mL of 4% PFA at a flow rate of 5 mL/min. 4% PFA in 0.1M PBS, pH 7, was prepared from powder (Thermo Scientific Chemicals, Cat. No. A1131336). Brains were rapidly dissected and post-fixed in 4% PFA overnight at 4°C. The eyes were enucleated following perfusion, and the cornea removed before placing into 4% EM-grade PFA in 0.1M PBS for 25 minutes on a nutating shaker at room temperature and then rinsed with 0.1M PBS with 0.1% Triton X-100 for 3 x 10 minutes. The following day, brains were rinsed with 0.1M PBS, and stored in 0.1M PBS with 0.1% Sodium Azide. An orienting cut was made in the left cortex before brains were embedded in 4% low-melt agarose (Fisher, Cat. No. 16520100) and sectioned at 100 μm using a vibratome (VT1200S Leica Microsystems Inc., Buffalo Grove, IL) and sections spread across 12-well plates containing 0.1M PBS with 0.02% Sodium Azide. For mapping retinorecipient locations of MAFB^+^ RGCs, serial sections of the entire brain were collected and immediately mounted on slides and cleared on slides using the CUBIC method ^91^. Slides were prepped for immunohistochemistry using the same protocol outline below.

Floating brain sections were blocked in 0.1M PBS containing 0.1% Triton X-100, and 2% donkey serum for 1 hour at room temperature. Tissue was then placed into primary antibody solution in blocking solution (see key resources table for specific dilutions) and incubated for 72 hours on a shaker at 4°C. Tissue was rinsed using blocking solution overnight, followed by incubation with a cocktail of secondary antibodies (1:500, Alexa Fluor 488, 546, 647) in blocking solution for 72 hours on a shaker at 4°C. Tissue was then rinsed with 0.1M PBS and 0.1% Triton X-100 overnight.

For retinal flatmounts, when possible, orienting cuts were made to denote the ventral retina using scleral landmarks. The retina was then removed from the eye cup if necessary (dependent on whether tissue was used for electrophysiology). Tissue was blocked in 0.1M PBS containing 0.1% Triton X-100, and 2% donkey or goat serum for 1 hour at room temperature. Tissue was then placed into primary antibody solution in blocking solution (see key resources table for specific dilutions) and incubated for 72 hours on a shaker at 4°C. Tissue was rinsed using blocking solution overnight, followed by incubation with a cocktail of secondary antibodies (1:500, Alexa Fluor 450,488, 546, 647) in blocking solution for 72 hours on a shaker at 4°C. Tissue was then rinsed with 0.1M PBS and 0.1% Triton X-100 overnight. For whole retinal flatmounts, four relaxing cuts were made.

For retinal sections, the entire globe was placed in 7.5% sucrose solution in 0.1M PBS for 1 hour and then in 15% sucrose solution in 0.1M PBS overnight. The lens was removed and the eye cups were placed in a cryomold with Optimal Cutting Temperature media (Fisher Scientific, Cat. No. 22-046-511) and frozen using methylbutane and dry ice. Retinas were sectioned at 20 μm using a cryostat (Leica CM3050 S) and mounted onto glass slides (Fisher Scientific, Cat. No. 12-550-15). Slides were left at room temperature for 1 hour to help with adherence. Slide mounted retinal sections were washed for 30 minutes in 0.1M PBS to rehydrate and then tissue was blocked in 0.1M PBS containing 0.1% Triton X-100, and 2% donkey serum for 1 hour at room temperature. Primary antibody solution in block was placed onto slides to incubate for 24 hours at 4°C (see key resources table for specific dilutions). Slides were rinsed using blocking buffer overnight, followed by incubation with a cocktail of secondary antibodies (1:500, Alexa Fluor 488, 546, 647) in blocking solution for 2 hours on a shaker at room temperature. Sections were washed with 0.1M PBS and counterstained with Hoechst 33342 (1:10,000, Life Technologies, Cat. No. H3570) for 10 minutes to visualize nuclei.

Rhesus macaque tissue was obtained from the Oregon National Primate Research Center. Tissue was fixed using 4% EM-grade PFA in 0.1M PBS for 30 minutes on a nutating shaker at room temperature and then rinsed with 0.1M PBS with 0.1% Triton X-100 for 3 x 10 minutes. Tissue was blocked in 0.1M PBS containing 0.1% Triton X-100, and 2% goat serum for 1 hour at room temperature. Tissue was then placed into primary antibody solution in blocking solution (see key resources table for specific dilutions) and incubated for 72 hours on a shaker at 4°C. Tissue was rinsed using blocking solution overnight, followed by incubation with a cocktail of secondary antibodies (1:500, Alexa Fluor 450,488, 546, 647) in blocking solution for 72 hours on a shaker at 4°C. Tissue was then rinsed with 0.1M PBS and 0.1% Triton X-100 overnight.

For all tissue processed for immunohistochemistry, following secondary antibody incubation, tissue was washed with 0.1M PB, and mounted onto glass slides using Fluoromount-G mounting medium (Fisher Scientific, Cat. No. OB100-01), coverslipped using #1.5 coverslips (Fisher Scientific, Cat. No. 12-541-026), and sealed using nail polish.

### RNAscope tissue preparation and assay

The Advance Cell Diagnostics (ACD) RNAscope Multiplex Fluorescent v2 assay (Advance Cell Diagnostics Inc., Cat.No. 323110) was adapted and modified for retinal flatmounts. For RNAscope, animals were deeply anesthetized using CO_2_ and intracardially perfused with 25 mL of 0.1M PBS with heparin (10 units/mL, Fisher Scientific, Cat. No. BP2524-50) followed by 25 mL of 4% PFA at a flow rate of 5 mL/min. The eyes were enucleated following perfusion, and the cornea and lens removed before placing into 4% EM-grade PFA in 0.1M PBS for 15 minutes. The retina was then isolated from the eyecup and attached photoreceptor side down onto a nitrocellulose membrane and fixed for an additional 30 minutes. Following fixation, the retina underwent a 3% H_2_O_2_ pretreatment for 20 minutes at room temperature. The retina was then mounted ganglion cell side up onto glass slides, dipped into dH_2_O and allowed to air dry for 1 hour. Slides underwent a EtOH dehydration series (50%, 70%, 100%), and washed 4 x 5 minutes with dH_2_O. Tissue was then permeabilized with RNAscope Protease IV for 30 minutes at 40°C, and washed 4 x 2 minutes with dH_2_O. Probes were warmed for 10 minutes at 40°C and diluted in diluent provided in ACD RNAscope Multiplex Fluorescent v2 kit, then added onto slides for 2 hours at 40°C. Slides were washed 4 times with 1x wash buffer. AMP 1 was added for 30 minutes at 40°C and then washed 4 times with 1x wash buffer. The same step was repeated for AMP 2. AMP 3 was added for 15 minutes then washed 4 times with 1x wash buffer. HRP steps are linked to probe channels. Probe channels were developed for 15 minutes at 40°C then washed 4 times with 1x wash buffer. Probe channels were stained using preproperate TSA Vivid or Opal fluorophores for 30 minutes at 40°C then washed 4 times with 1x wash buffer. HRP blocker was added for 30 minutes at 40°C then washed 4 times with 1x wash buffer. For immunohistochemistry following RNAscope, tissue was blocked for 10 minutes in blocking solution, incubated with appropriate primary antibodies in block solution for 1 hour at 40°C then washed with 0.1M PB 4 x 10 minutes. Secondary antibodies in block solution along with Hoechst 33342 (1:10,000, Life Technologies, Cat. No. H3570) was added for 30 minutes at 40°C, washed with 0.1M PB 4 x 10 minutes. Tissue was covered with Fluoromount-G mounting medium (Fisher Scientific, Cat. No. OB100-01), coverslipped using #1.5 coverslips (Fisher Scientific, Cat. No. 12-541-026), and sealed using nail polish.

### Fluorescence image acquisition

Imaging was performed on either a Zeiss Axio Imager M2 upright microscope equipped with an Apotome.2 or a Leica TCS SP8 laser scanning confocal microscope. Axio Imager M2 images were captured using a 20x/0.8 NA air objective and acquired using the Zeiss Zen Imaging software. Leica TCS SP8 laser scanning confocal images were acquired using a 40x/1.3 NA oil immersion objective and the Leica Application Suite X (LAS X) software. Z-stack images were acquired and analyzed offline in ImageJ/FIJI ^92^. For imaging of MAFB^+^ RGC termination in the brain, slides were imaged at 5x using a Zeiss Axioscan7 slide scanner by the OHSU Advanced Light Imaging Core (RRID: SCR_009961). Brightness and contrast were adjusted in ImageJ/FIJI (Version 2.14.0/1.54f) to improve visibility of images for display. Figures were composed in Affinity Designer 2 (2.2.0).

### Quantification and statistical analysis

#### Image quantification

For mapping retinorecipient locations, serial brain sections from four animals were imaged for fluorescence detection of GFP, 647-CTB and Hoechst. For retinal flatmount imaging, 2-3 images per retina were acquired from various locations within the retina (central, middle, and periphery), averaged, and at least two animals from different litters were used for analysis. For sparsely labeled RGCs, 1-5 images per retina (dependent on density of viral labeling) were acquired, avoiding cells in the far periphery of the retina. For retinal section imaging, 3 images, each from different sections, were acquired from regions of the retina closest to the optic nerve. For image processing, background subtraction (rolling ball = 50), despeckle, and a median filter (radius = 2) were applied. At least two animals from different litters were used for analysis.

### Data and code availability

R scripts were written with the assistance of ChatGPT-4 to analyze morphological, retinorecipient location, and electrophysiological data. All code was refined and verified by NL. R (R version 4.3.2), and Rstudio (Posit Software, Version 2024.12.1^+^563), was used to further analyze the data and generate all graphs.

### Statistics

Statistical analyses were performed using R. To test for differences across age groups, a two-sided one-way ANOVA with α = 0.05 was applied. For morphology and physiology data, normality was assessed within each type using the Shapiro-Wilk test. For normally distributed data, homogeneity of variance was assessed using Bartlett’s test. Normally distributed variables with unequal variances were analyzed using Welch’s ANOVA followed by Games-Howell pairwise comparisons. Variables that did not satisfy the normality assumption were analyzed using the Kruskal-Wallis test followed by Dunn’s pairwise comparisons with Benjamini-Hochberg correction. Comparisons between two independent groups were performed using Student’s t-test, Welch’s t-test, or the Wilcoxon rank-sum test, as appropriate. Paired ON and OFF measurements from the same cells were compared using two-sided paired Wilcoxon signed-rank tests. All tests were two-sided, and statistical significance was defined as α = 0.05. Sample sizes (n) represent the number of cells included for each analysis reported in the corresponding figure legends or results tables. No statistical methods were used to predetermine sample sizes.

### Single cell transcriptome mining

To re-analyze MAFB^+^ clusters, we acquired single-cell transcriptomic data from the RGC atlas generated by Tran et al., 2019, publicly available through GEO (accession GSE137400). After standard quality control, keeping genes expressed in more than three cells and cells with over 200 detected genes, we integrated the data across samples (aRGC1-9) and batches (1-3) using the Seurat v4 pipeline to reduce technical variability ^93^. Normalization was performed with SCTransform(), with mitochondrial and ribosomal expression regressed out, and 3,000 highly variable genes were used as integration anchors. From the integrated dataset, MAFB^+^ clusters were isolated using published annotations, followed by dimensionality reduction using the optimal number of principal components (PCs) based on the scree plot elbow (9 PCs), UMAP visualization, and differential expression analysis to resolve MAFB^+^ clusters.

A custom R script was written to perform differential gene expressed genesion analysis among selected *Mafb*^+^ clusters were identified from the SCTransform-normalized assay using the Seruat FindAllMarker() function. PrepSCTFindMarkers() was applied to recorrect the SCT counts using a common sequencing-depth reference across the SCT models. Each cluster was compared with all remaining cells using a two-sided Wilcoxon rank-sum test. Testing was limited to genes detected in at least 25% of cells in either comparison group and exhibiting an average log2 fold change of at least 0.25. Only positively enriched genes were retained. P-values were adjusted across genes using the Bonferroni procedure implemented by Seruat, and genes with adjusted p < 0.05 were considered significant. using a one-versus-all comparison strategy. For each cluster of interest, cells assigned to that cluster were compared against all cells belonging to the remaining selected clusters. Cell identities were determined using cluster annotations from Tran et al., 2019, and gene expression matrices were subset accordingly. Differential expression for each gene was assessed using row-wise t-tests between the target cluster and the pooled comparison group. Genes were ranked by absolute t-statistic to identify those exhibiting the strongest differential expression, and highly ranked genes were used to identify candidate molecular markers distinguishing each cluster. Click or tap here to enter text.

### MAFB^+^ predicted immunohistochemistry markers

The predicted frequency (p_f_) percentage for each immunohistochemical marker was calculated by first identifying all *Mafb*^+^ t-types expressing the marker (scaled mean expression from scRNAseq must be greater than or equal to 0.30). The predicted frequencies of these t-types were summed and divided by the total relative frequency of *Mafb*^+^ t-types, then multiplied by 100 to get a percentage.

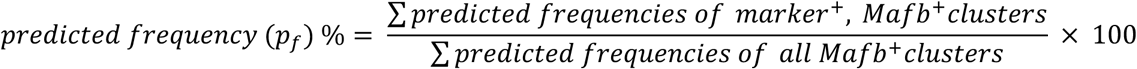

To calculate the observed frequency (o_f_) percentage for each marker, the total numer of mCherry^+^ cells within a 300 x 300 μm region of interest was manually counted using the Cell Counter plug-in from ImageJ/Fiji. Cells that colocalized with the marker of interest and mCherry were manually counted in the same region of interest. Colocalized cells were divided by the total number of mCherry cells, then multiplied by 100 to get a percentage.

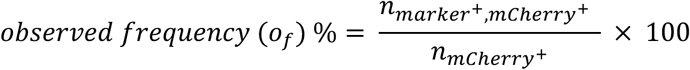

### MAFB^+^ RGC morphological analysis

To identify IPL stratification depth, the rectangle tool in ImageJ/FIJI was used to draw a 200 x 200 pixel box (5.2826 pixels per μm) in the dendritic tree and the z-plot profile analyzed for fluorescence intensity of each channel. The distance axis for each profile was normalized from 0 to 100%. Signal intensity within each channel was subsequently normalized from 0 to 1 using the minimum and maximum intensity measured within that profile. The ChAT/VAChT intensity profile was smoothed using a three-point moving average. Local maxima were identified from points where the smoothed intensity exceeded that of the immediately adjacent points. The two local maxima with the greatest amplitudes were selected as the major cholinergic bands and the more proximal of these two peaks (S4) was used as the alignment reference. For each profile, the distance axis of ChAT/VAChT, TH and GFP was shifted such that the S4 peak was assigned depth 0. The positions of the S2 and S1 boundaries were retained according to their relative locations within the normalized 0ñ100% IPL profile following alignment. TH labeling (S1) was plotted as an additional anatomical reference but was not used to determine the alignment or IPL-depth normalization. The GFP z-profile depth was then normalized to ChAT/VAChT plexus depthas reference points for S2 and S4 and/or TH staining for S1. Cells with the same stratification pattern were then averaged. Soma area was measured by using the freehand selection tool. To measure the convex hull of the RGC dendritic arbor, the polygon selection tool was used to connect the tips of the dendrites in each stratum and then convex hull was applied. For bistratified RGCs, a convex hull was drawn around the first dendritic plexus, closest to GCL, to obtain the dendritic field of plexus 1 (DF_1_) and around the second dendritic plexus to obtain dendritic field of plexus 2 (DF_2_). DF ratio was calculated using the following:

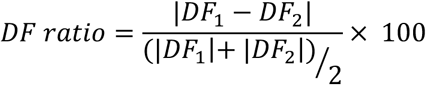

Dendritic skeletons were made using the ImageJ/FIJI plug-in Simple Neurite Tracer (SNT) ^39^. The following metrics were collected using SNT: z-depth, total branch length, total number of branches, and number of tips. Arbor density was calculated using (total branch length / dendritic field area) * 100. Arbor complexity was calculated using (total number of branches / total branch length) * 100. An ImageJ/FIJI macro was created to calculate SI. Using the binarized dendritic skeleton, where the background is black and dendrites are white, the soma of the cell was manually indicated, and the longest dendritic tip was automatically identified by finding the furthest white pixel from the soma. A circle was drawn to encompass all dendritic branches and then split into 12 equal segments. Each segment was analyzed to find the number of white pixels within the area (*S_i_*). Individual were compared to the average number of white pixels for the other slices (*S_other_*). to obtain a slice symmetry index (*SSI*). The average number of white pixels for *S_other_* is calculated by summing the of all other slices and dividing it by 11. To obtain the average SI for a given cell, all slice symmetry indexes are summed and divided by 12. The average SI was reported.

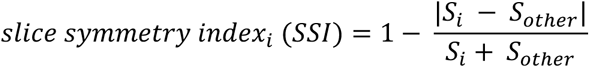

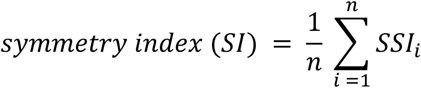

### Unbiased morphometric clustering

Morphometric clustering was performed using quantitative measurements obtained from reconstruction MAFB^+^ RGCs. Four feature sets were evaluated independently, each consisting of different combinations of morphological parameters. Morphological features were centered and scaled prior to PC analysis using the prcomp() function in R. The minimum number of PCs required to explain at least 80% of the total variance was retained. The optimal number of clusters (k) was determined using gap statistics (clusGap() function in R) and k chosen based on firstSEmax. To visualize relationship between RGCs, two-dimensional embedding plots were generated using tSNE applied to the retained PC scores. A perplexity of 10 was used, and random seed were fixed for PCA, clustering and tSNE for reproducibility.

To determine how Neurobiotin-filled MAFB^+^ f-types were distributed within morphological groupings, recorded cells were incorporated directly into the clustering analysis. Morphological measurements from Neurobiotin-filled MAFB^+^ f-types were extracted using the same parameters employed for Supplementary Figure 7H feature set. The cells were combined with the reconstruction dataset and underwent identical preprocessing, dimensionality reduction, gap stat analysis, k-means clustering and tSNE visualization.

### Electrophysiology data analysis

A custom R script was written to extract spike times, classify the polarity and kinetics of cells, and analyze feature selectivity of RGCs. Current traces were high-pass filtered at 0.2 Hz using a second-order Butterworth filter and low-pass filtered at 700 Hz using a fourth-order Butterworth filter. The filtered traces were differentiated, and spikes were identified when the first derivative exceeded 50% of the maximum derivative observed within each sweep. To prevent multiple detections of the same spike, a refractory period of 2 ms was enforced before subsequent spikes could be detected. Spike frequencies were calculated in 50-ms bins and averaged across all stimulus repetitions. Spike detections were visually verified in representative recordings by overlaying automatically detected events on individual current traces for each sweep. Waveforms surrounding detected events were also extracted and overlaid to confirm consistent spike-like morphology. The same detection parameters were then applied uniformly across recordings. Recordings were excluded if excessive baseline noise prevented reliable spike detection following visual inspection prior to analysis or if no light-evoked responses were observed to any of the visual stimuli (blue, green, or black). The following standard metrics were identified using three trials that each contained five sweeps of green/blue and black light step stimuli.

*Baseline firing rate*: mean firing rate before spot presentation across all trials.

*Peak firing rate*: highest firing rate (baseline subtracted) in 50 ms bins at light onset or offset.

*Latency to peak*: time from light onset or offset until the peak firing rate.

*Final firing rate*: mean firing rate halfway through the presentation of a stimulus.

*Final to peak (FP) index*: (final firing rate - baseline firing rate) / (peak firing rate - baseline firing rate)

After the standard metrics were computed, cells were then classified as ON, OFF, or ON-OFF based on the presence or absence of firing during the presentation of the light step stimuli. Cells were then classified as transient or sustained responders based on their FP index. Cells with an FP index greater than or equal to 0.15 were classified as sustained. Any cells that did not pass the initial polarity classification were omitted from any further analysis. To determine the receptive field size of MAFB^+^ RGCs, spots with diameters ranging from 25-1200 μm were presented on the RF center. The diameter that elicited the highest spike frequency was deemed the cell’s RF size. For contrast, the contrast that provoked the highest spike frequency rate was determined to be the cell’s preferred contrast. Orientation selectivity index (OSI) and preferred orientation angle were calculated based on Nath & Schwartz, 2016. OSI was calculated using the vector sum for angles. All angles are doubled, to allow angle vectors that are opposite from one another (i.e. 30° and 210°) to point in the same orientation. Vectors that align are then stacked. OSI values range from 0 (not orientation-selective) to 1 (highly orientation-selective), with the cut off for a cell being orientation selective set at OSI ≥ 0.33.

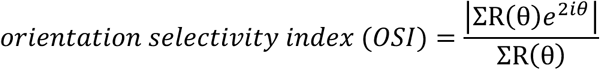

Direction selectivity index (DSI) was calculated based on methods provided in Duan et al., 2014. For each direction, spike frequencies were averaged across 2-3 repeated trials to obtain an average response magnitude. DSI was calculated as the magnitude of the vector sum of direction-specific responses divided by the sum of the response magnitudes across all directions. DSI values range from 0 (not direction selective) to 1 (highly direction selective), with the cut off for a cell being direction selective set at DSI ≥ 0.33.

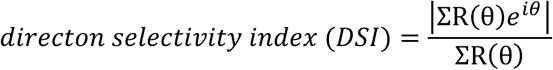

## Data availability

The data that support the findings of this study are available from the corresponding author upon reasonable request.

## Acknowledgements

We would like to thank the members of the Wright lab, Sivyer lab, and Arpiar Saunders for their feedback on the manuscript. We would like to thank the Oregon National Primate Research Center for the macaque tissue and the OHSU Advanced Light Imaging Core for imaging help and acquisition. This work was supported by the National Institute of Health grants: R01EY032057, R01EY032564, T32NS07466, T32EYE023211, and P30EY010572.

## Contributions

N.L., B.S., and K.M.W. designed the research. O.H. performed the experiments and analysis in Figure 1D, F and Supplementary Figure 2C-E with supervision from N.L. and K.M.W. N.L. performed the rest of the experiments. Analysis of data was done by N.L. with help from B.S. and K.M.W. The manuscript was written by N.L. and K.M.W. with input from B.S. Funding for this project was secured by K.M.W. and B.S, and maintained by J.L.

## Corresponding Author

Correspondence to Kevin M Wright.

---

**Supplementary Figure 1.**
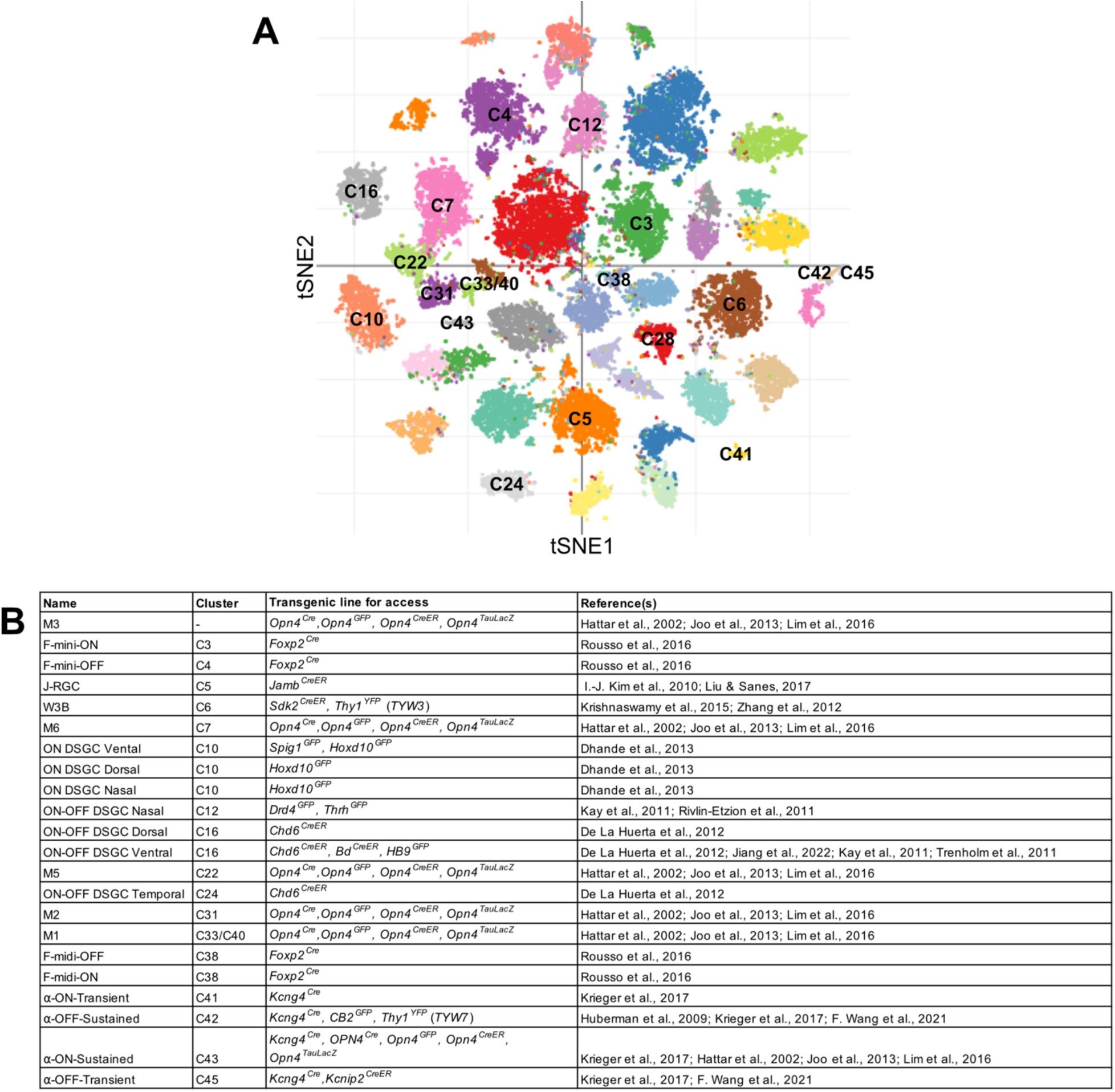
Known RGC subtypes and available transgenic mouse lines for labeling and experimental manipulations. **(A)** tSNE plot showing transcriptomically defined RGC clusters in the adult mouse retina. Clusters corresponding to RGC types with genetic access are annotated. Adapted from Tran et al., 2019**. (B)** Table listing the identified RGC types, their corresponding transcriptomic cluster assignment, and transgenic mouse lines used for selective labeling and manipulation.

**Supplementary Figure 2.**
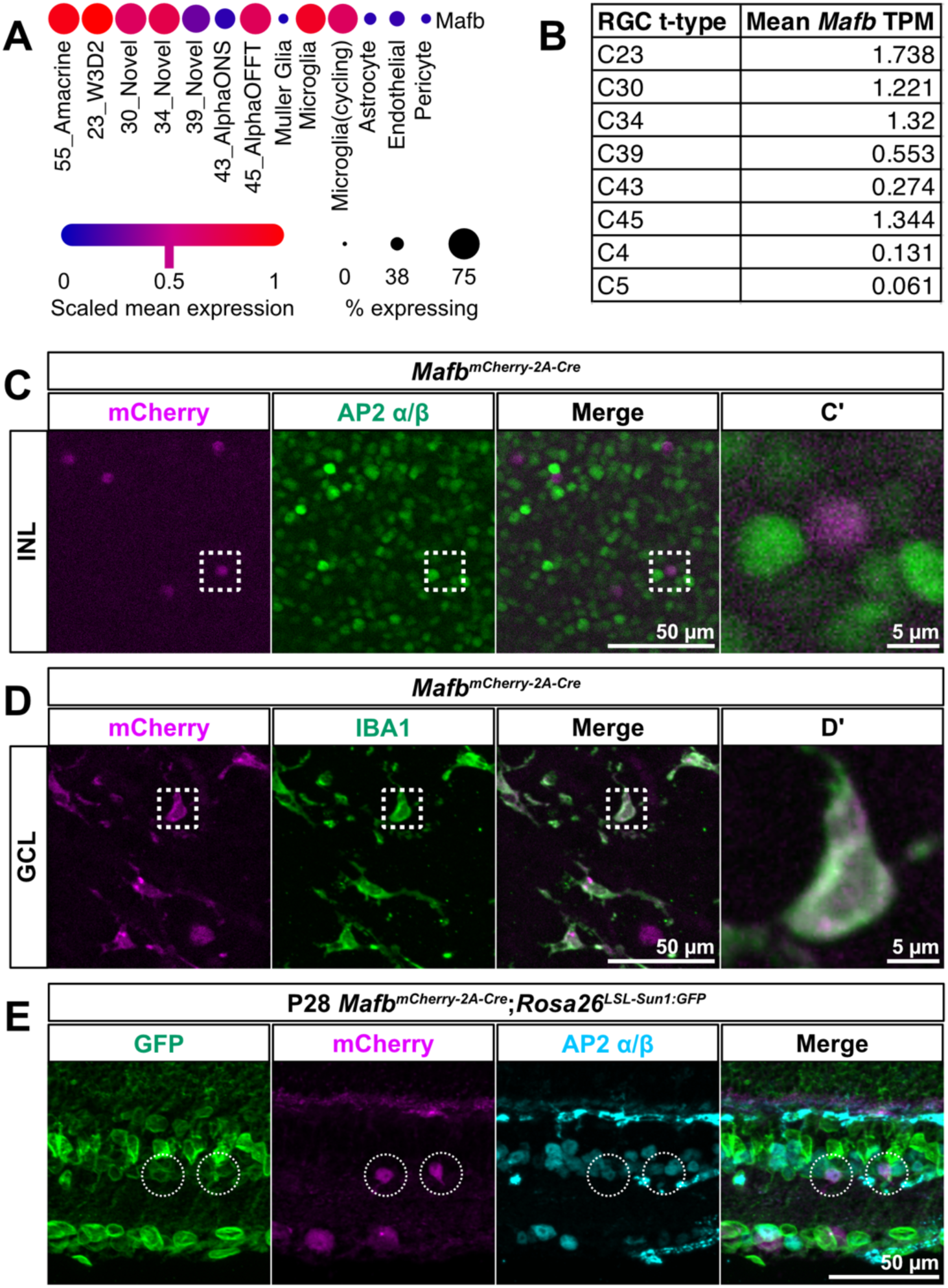
*Mafb* is expressed in other retinal neurons. **(A)** *Mafb* expression in retinal cells, dot plot adapted from MRCA atlas ^9^. **(B)** Normalized mean *Mafb* transcripts per million (TPM) for *Mafb*^+^ t-types (C23, C30, C34, C39, C43, and C45) and two *Mafb*^-^ t-types (C4 and C5) from Tran et al., 2019. **(C)** *En face Mafb^mCherry-2A-Cre^* retina immunostained for mCherry (magenta) and AP2α/*β* (green) show colocalization in the INL. **(C’)** Magnified view of the white outlined box in composite image **(D)** *En face Mafb^mCherry-2A-Cre^* retina stained with mCherry (magenta) and pan-microglia marker IBA1 (green) show colocalization in the GCL. **(D’)** Magnified view of the white outlined box in composite image. **(E)** Retinal section from P28 *Mafb^mCherry-2A-Cre^*;*Rosa26^LSL-Sun1:GFP^* immunostained for GFP (green), mCherry (magenta) and pan-AC marker, AP2α/β (cyan). Circled areas show mCherry^+^ ACs with sustained expression of *Mafb*. Scale bar = 50 μm in **(C)**, **(D)**, and **(E)**. Scale bar = 5 μm in **(C’)** and **(D’)**.

**Supplementary Figure 3.**
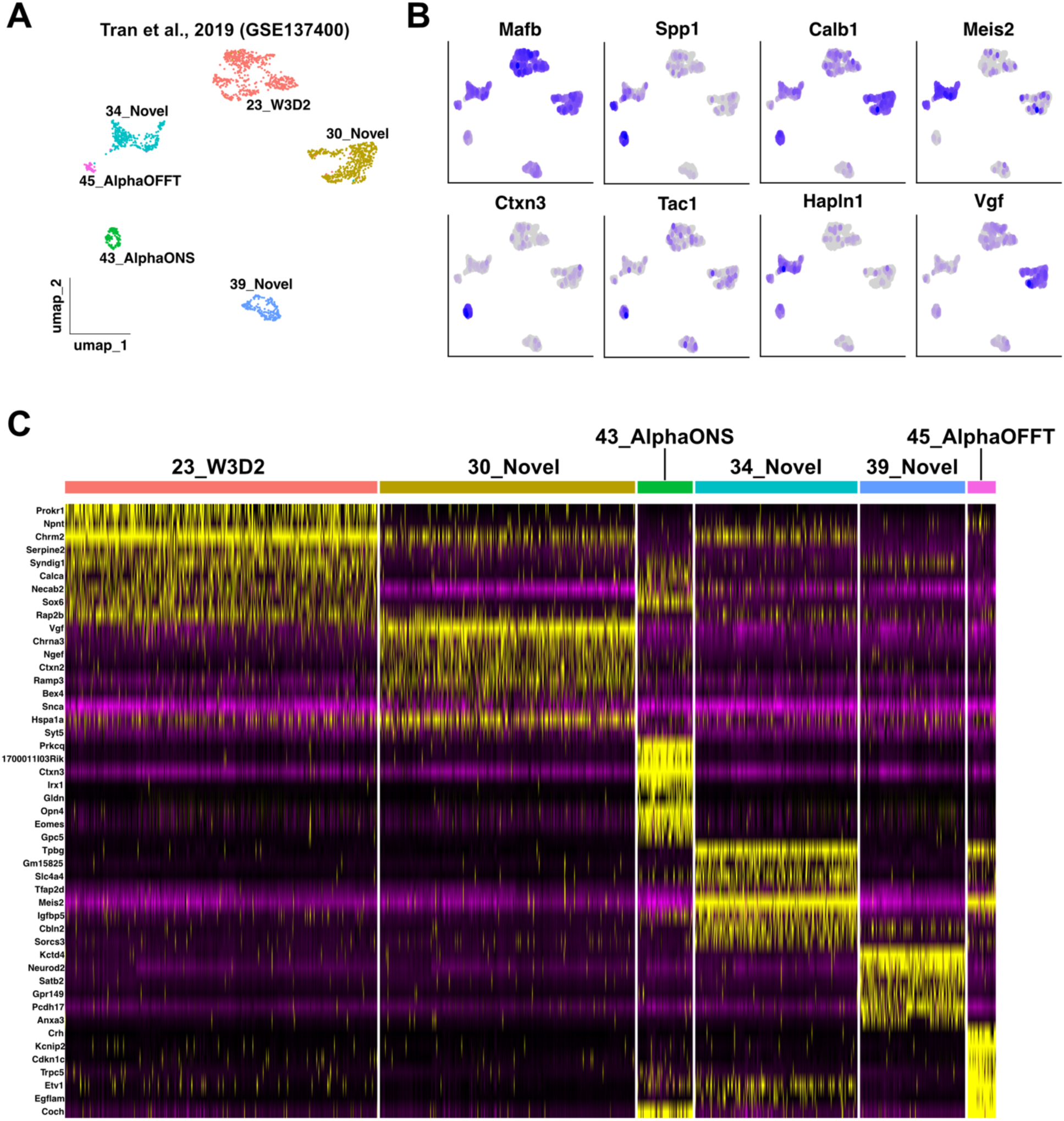
Reclustering and gene expression of *Mafb*-expressing RGCs. **(A)** 2D UMAP embedding of reclustered *Mafb*-expressing RGCs from Tran et al., 2019. **(B)** Candidate cluster-enriched molecular markers. **(C)** Heatmap showing differentially expressed genes across clusters, ordered by cells as columns and genes as rows.

**Supplementary Figure 4.**
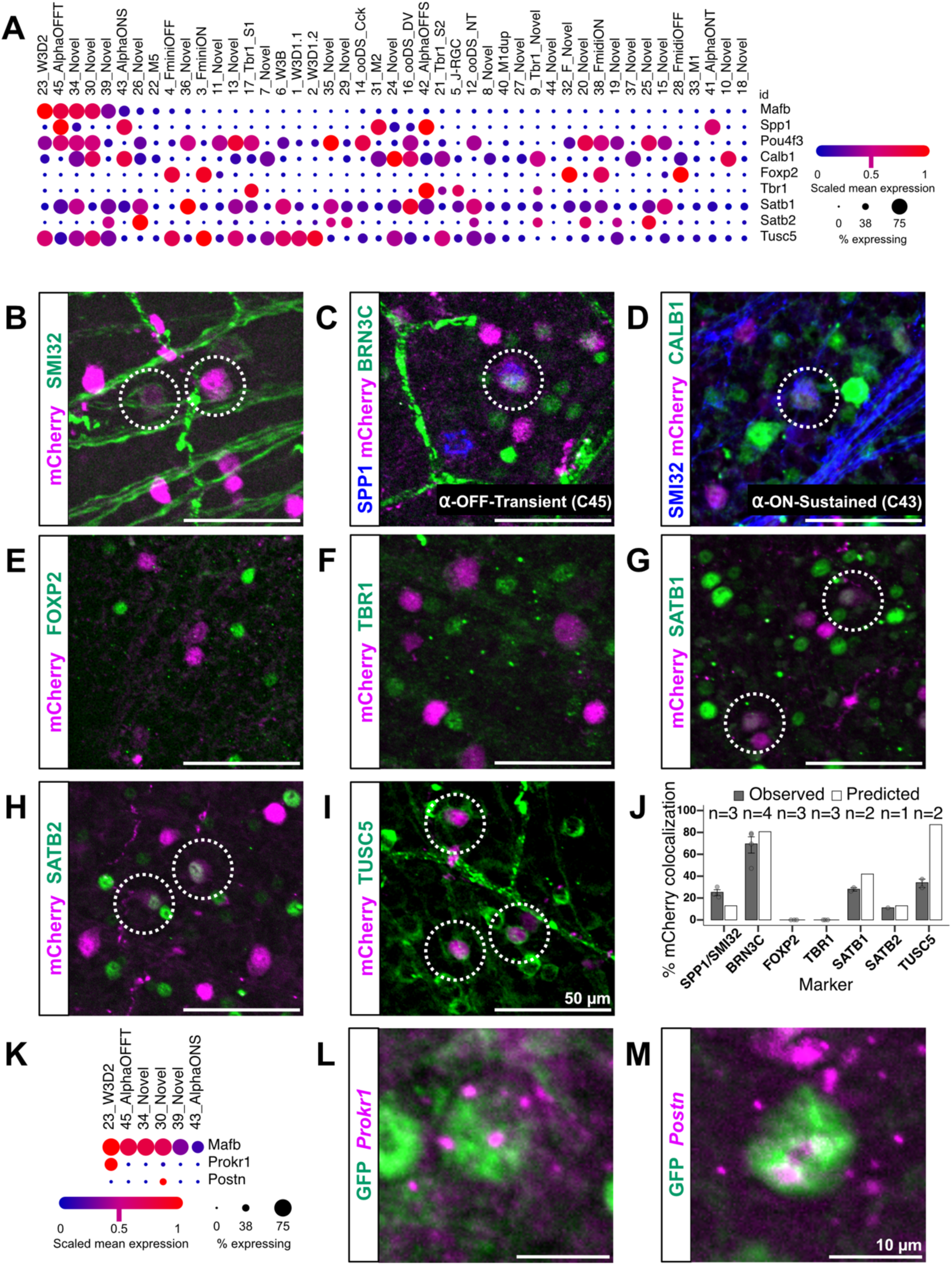
Molecular identification of MAFB^+^ RGCs. **(A)** scRNAseq dot plot showing expression of *Mafb*, *Spp1*, *Brn3c* (*Pou4f3*), *Calb1*, *Foxp2*, *Tbr1*, *Satb1*, *Satb2*, and *Tusc5*. **(B-I)** *Mafb^mCherry-2A-Cre^* retinas immunostained for mCherry (magenta) and various markers (green/blue) to identify molecular subtypes. Scale bar = 50 μm. Dotted circles indicate colocalization. **(B)** SMI32, a marker for α-RGCs. **(C)** Triple labeled SPP1^+^ (blue), BRN3C^+^ (green), mCherry^+^ cells are α-OFF-Transient RGCs (C45). **(D)** Triple labeled SMI32^+^ (blue), CALB1^+^ (green), mCherry^+^ cells are α-ON-Sustained RGCs (C43). **(E)** FOXP2, a marker for F-RGCs. No colocalization seen between FOXP2 and mCherry. **(F)** TBR1, a marker for TBR1^+^ RGCs. No colocalization seen between TBR1 and mCherry. **(G)** SATB1, a marker for DSGCs. **(H)** SATB2, a marker for other novel RGC types. **(I)** TUSC5, a marker for T5-RGCs. **(J)** Percent colocalization of indicated markers along with the predicted colocalization based on scRNAseq (see methods for detailed calculations). Values are presented as (marker, predicted labeled *Mafb*^+^ t-types if applicable, predicted frequency (p_f_), observed frequency (o_f_) ± SEM, N = animals). SPP1/SMI32, C43/C45, p_f_ =12.90%, o_f_ = 25.25% ± 2.67%, N = 3 animals; BRN3C, C23/C30/C34/C45, p_f_ =80.65%, o_f_ = 69.52% ± 6.48%, N = 4 animals; FOXP2, p_f_ =0%, o_f_ = 0%, n = 3; TBR1, pf =0%, o_f_ = 0%, N = 3 animals; SATB1, C34/C39/C45, p_f_ =41.94%, o_f_ = 28.08% ± 1.47%, N = 2 animals; SATB2, C39, p_f_ =12.90%, o_f_ = 11.10%, N = 1 animals; TUSC5, C23/C30/C34/C39, p_f_ =87.10%, o_f_ = 34.04% ± 3.27%, N = 2 animals. **(K)** scRNAseq dot plot showing selective expression of *Prokr1* and *Postn* in C23 and C30. **(L-M)** RNAscope for *Prokr1* **(L)** and *Postn* **(M)**. Scale bar = 10 μm.

**Supplementary Figure 5.**
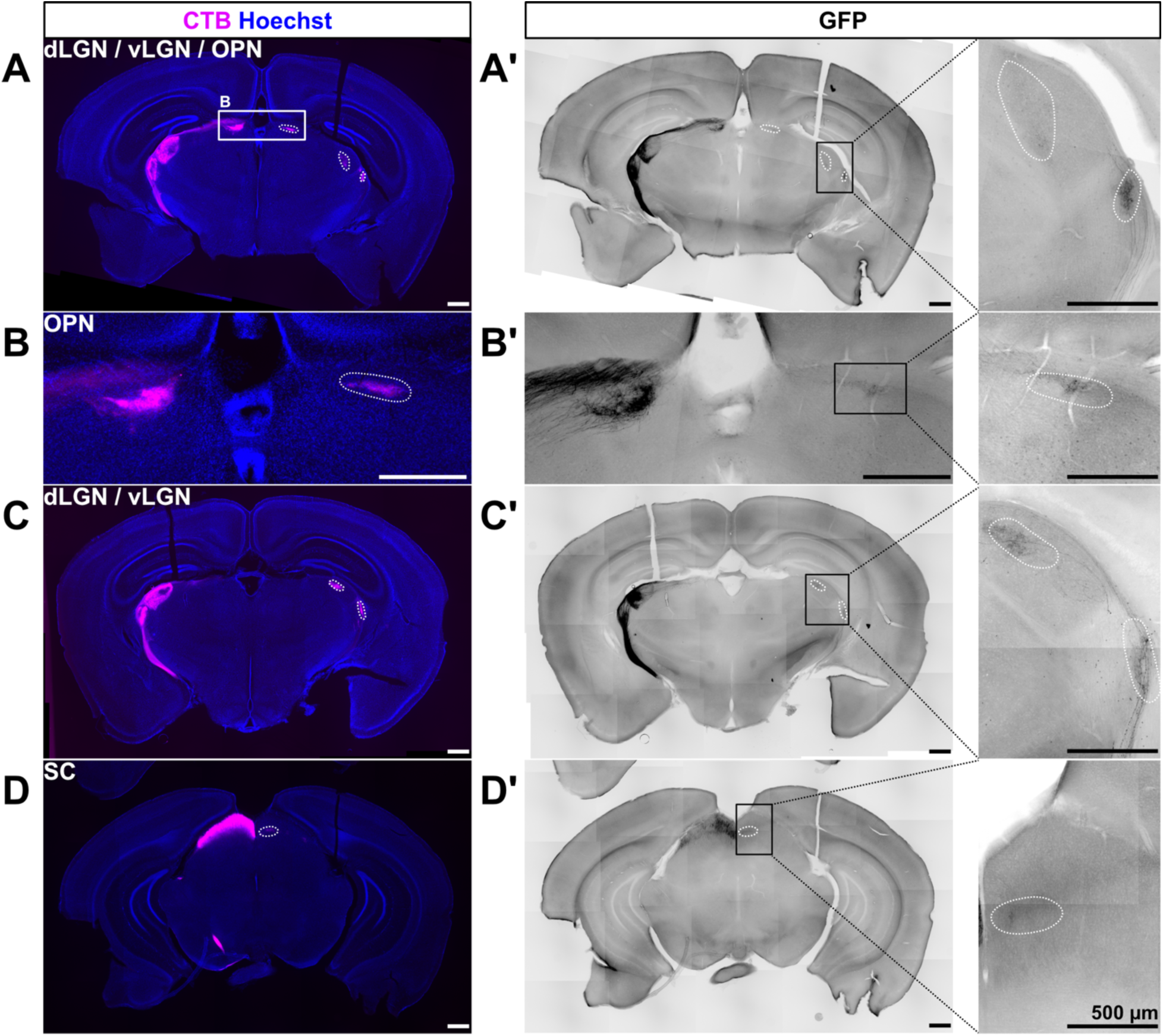
Contralateral and ipsilateral projection pattern of MAFB^+^ RGCs. High titer *AAV2-CAG-FLEX-EGFP* was IVT injected unilaterally into *Mafb^mCherry-2A-Cre^* mice to map contralateral and ipsilateral projections of MAFB^+^ RGCs to retinorecipient brain regions. Cholera toxin B (CTB) conjugated to Alexa Fluor 647 (magenta) was IVT injected unilaterally into the same eye and used as an anterograde tracer to label all retinal projections to retinorecipient brain regions. **(A-D)** Coronal brain sections showing CTB labeling of retinal projections (magenta). Boxed area in panel **A** corresponds to inset **B**. Scale bar = 500 μm. **(A’-D’)** Corresponding brain sections showing GFP-labeled axons. Dotted lines indicate ipsilateral retinorecipient location. Boxed areas indicate magnified view of dotted regions. Sparse GFP^+^ terminals were seen in **(A’, C’)** vLGN, **(B’)** OPN and **(A’, C’)** dLGN. No obvious termination is seen in the **(D’)** ipsilateral SC.

**Supplementary Figure 6.**
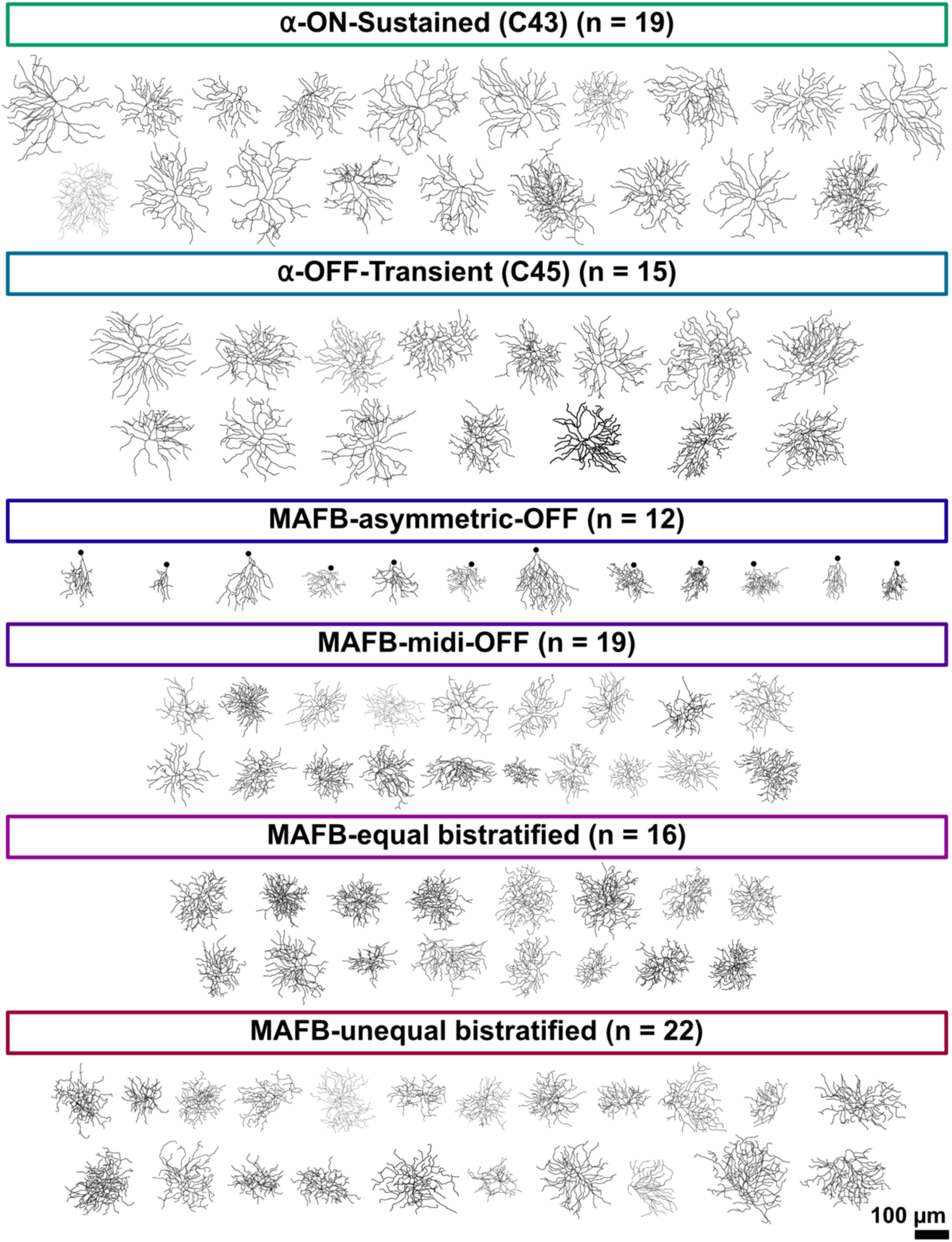
Dendritic skeletons of reconstructed MAFB^+^ RGCs. All reconstructed MAFB^+^ RGC dendritic skeletons separated by m-type, N = 55 animals. RGCs were reconstructed using ImageJ/FIJI Simple Neurite Tracer. Skeletons were dilated for visibility. Scale bar = 100 μm.

**Supplementary Figure 7.**
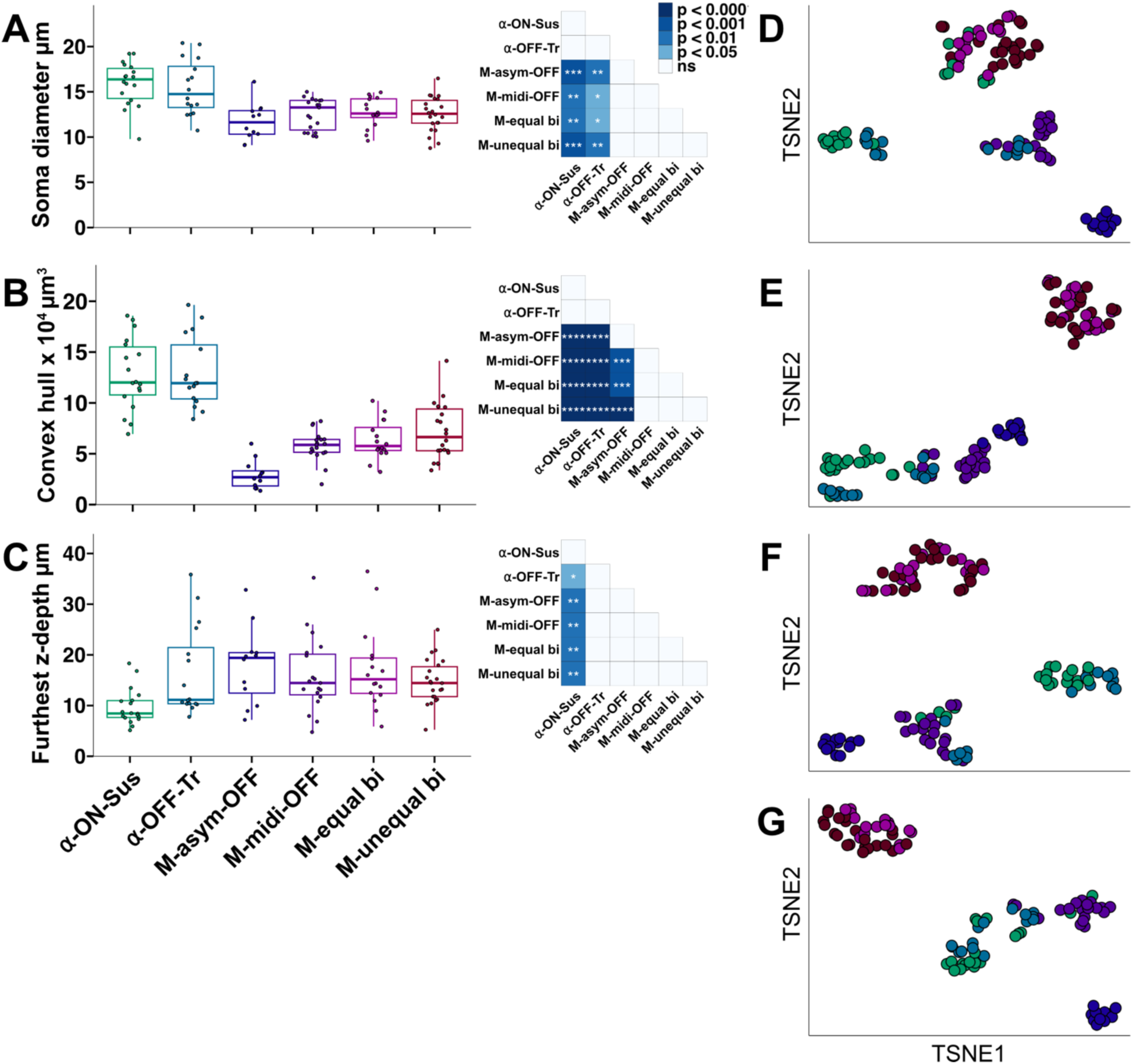
Additional morphometric analyses of MAFB^+^ m-types. **(A-C)** Quantification of **(A)** Soma diameter, **(B)** Convex hull, **(C)** Furthest z-depth. Each point represents an individual reconstructed cell, with boxplot indicating median and interquartile range. A Kruskal-Wallis test, *df* = 5, and post hoc Dunn’s test with Benjamini-Hochberg correction was performed for statistical analysis of **(A)** and **(C)**. A Welch’s ANOVA, *df* = 5, and post hoc Games-Howell test was performed for statistical analysis of **(B)**. Heatmaps summarise pairwise statistical comparisons between MAFB^+^ RGC m-types. **(D-G)** tSNE plot using different morphometric parameters and number of principal components (PCs). Color corresponds to m-type based on **A-C**. **(D)** 4 PCs retained, k = 5, consisting of the following metrics: soma diameter, discrete IPL stratification, bistratifed DF ratio, convex hull, SI, and terminal tips. **(E)** 3 PCs retained, k = 4, consisting of the following metrics: soma diameter, discrete IPL stratification, DF_1_ and DF_2_ diameter, SI, and terminal tips. **(F)** 3 PCs retained, k = 5, consisting of the following metrics: soma diameter, IPL stratification depth, DF_1_ and DF_2_ diameter, and SI. **(G)** 4 PCs retained, k = 6, consisting of the following metrics: soma diameter, discrete IPL stratification, monostratified or bistratified, bistratifed DF ratio, convex hull, SI, total branch length, and total number of branches.

**Supplementary Figure 8.**
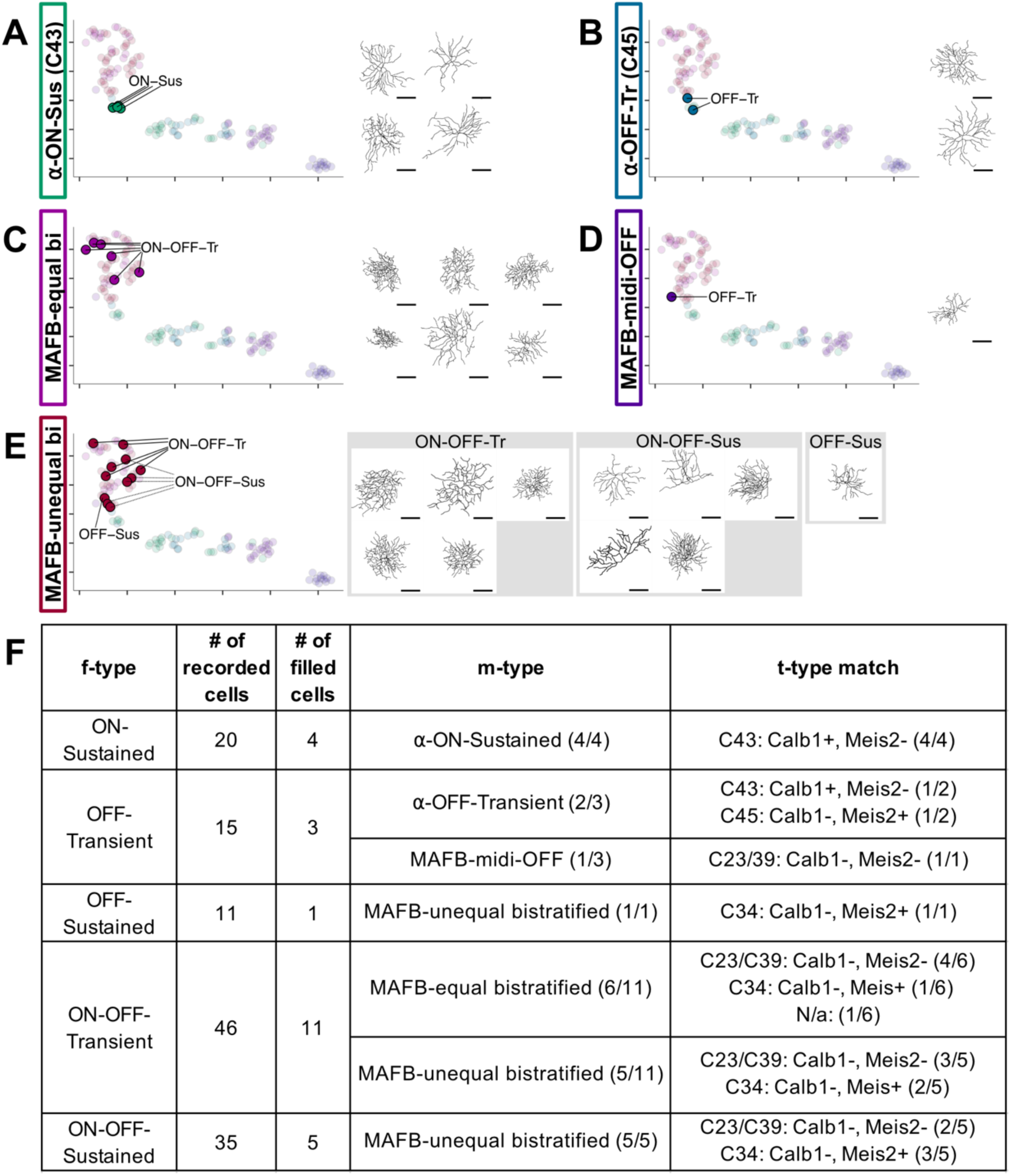
Unbiased clustering of f-types based on m-type. **(A-E)** tSNE plot with integration of m-types from recorded RGCs. Representative cells from each cluster are highlighted and annotated with their functional response. Reconstructions to the right show example dendritic morphologies corresponding to each functional group. **(F)** Summary table with number of recorded cells per f-type, number of filled cells, t-type match, and m-type match.

**Supplementary Figure 9.**
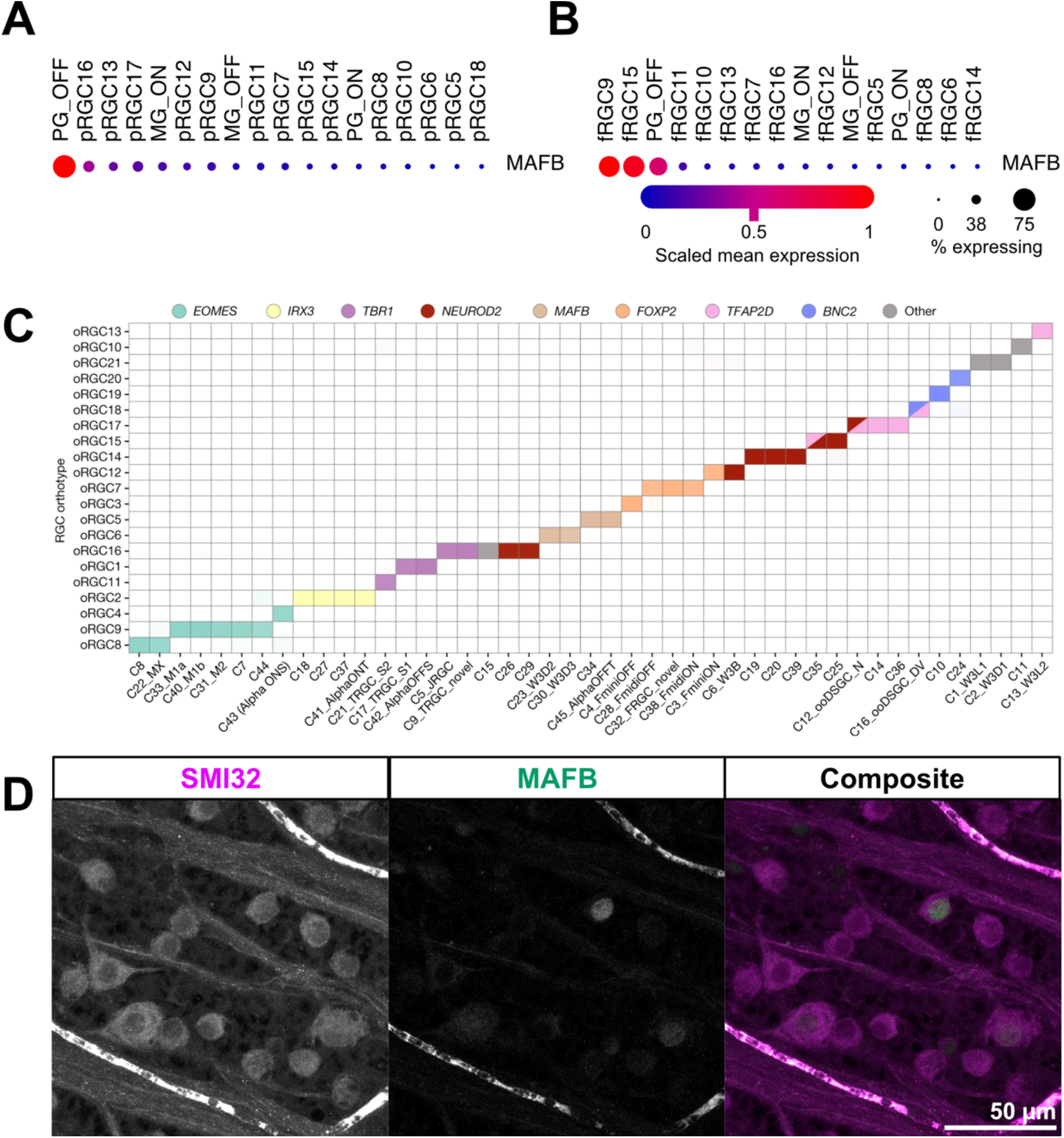
*Mafb* is expressed in nonhuman primate RGCs. **(A)** Dot plot showing *Mafb* expression in peripheral macaque RGCs. **(B)** Dot plot showing *Mafb* expression in foveal macaque RGCs. **(C)** Orthotypes of RGCs and mapping of respective mouse RGC clusters. Color denotes transcription factor identity. Panel from Hahn et al., 2023. **(D)** Macaque retinal flatmount from left eye, superior temporal region stained with SMI32 (magenta) and MAFB (green). Scale bar = 50 μm.

